# Single-cell tracking of genetically minimized *Salmonella* reveals mechanisms of effector gene cooperation

**DOI:** 10.1101/2025.05.12.653496

**Authors:** Wesley B. Burford, Hrag Dilabazian, Laura T. Alto, Duo Ma, Angela B. Mobley, Arun Radhakrishnan, J. David Farrar, Neal M. Alto

## Abstract

Bacterial pathogens encode secretory systems that deliver large repertoires of effector proteins directly into host cells. While many effector proteins have been characterized biochemically, it is unclear where and when they function within complex cellular systems of host tissues. This problem exists because of the extensive genetic and functional redundancies found in large effector protein repertoires. Here, we coupled targeted genome minimization with single cell mass cytometry to track the cellular location where *Salmonella* Typhimurium (*S.*Tm) SPI-2 Type Three Secretion System effector proteins function in a time-resolved animal model of infection. This approach revealed the temporal progression of *S.*Tm transmission within a complex tissue and pinpointed effector genes responsible for pathogen dissemination between host immune cell types. We further show how coupling two effector gene networks acquired during distinct episodes of bacterial evolution has shaped the cellular and tissue tropism of non-Typhoidal *Salmonella* species. Together, these results illustrate a top-down genetic approach to interrogate host-pathogen interactions hidden by functional redundancies within large virulence factor gene networks.

## Introduction

Gram-negative bacterial pathogens encode dedicated protein secretory systems (e.g. T3SS, T4SS, and T6SS) that translocate “effector” proteins into the host cell cytoplasm^1^. Many pathogens have acquired very large repertoires of effector genes, such as *Legionella pneumophila* that delivers over 300 effector proteins into host amoeba and human alveolar macrophage^2^. While the secretory systems are essential for survival of bacteria within the host organism, previous studies have shown that knocking out individual effector proteins often have little effect on virulence^3^. It has been suggested that large effector gene repertoires are fraught with genetic and functional redundancies that mask the essential nature of individual effector genes. Unfortunately, these redundancies have complicated genotype to phenotype relationships that are the cornerstone of forward- and reverse-genetic approaches used to characterize effector protein activities within the host.

Recent advances in the field of synthetic biology have presented a potential solution to the problem of functional redundancy in host-pathogen interactions. That is, fundamental principles of pathogenesis may be elucidated through a process known as “genome minimization,” which involves the experimental reduction of an organism’s genetic material while maintaining essential functions^4–6^. To be more precise, genome minimization strives to preserve genes and sequences vital for the organism’s growth, reproduction, and response to environmental challenges, while discarding non-essential or redundant elements. The quest for a minimal effector genome in a mammalian pathogen has gained momentum over the past decade, due in large part to a handful of pioneering studies^7–9^. Recently, we minimized the effector gene repertoire translocated by the *Salmonella* Pathogenicity Island 2 (SPI-2) Type 3 Secretion System T3SS^9^. Using phenotypic benchmarks to eliminate non-essential SPI-2 effector genes, we successfully reduced the SPI-2 T3SS effector gene repertoire by over 80%. We engineered an *S.*Tm strain that secretes just five effector proteins – SopD2, SteA, SifA, SseF and SseG – and this small genetic network was found to be sufficient for *S.*Tm to survive and replicate to high levels within model epithelial cell and macrophage growth in culture^9^. While the molecular mechanisms of these effector proteins have been studied independently (reviewed in ^10,11^), our work suggested they function as a cohesive unit that allows *S*.Tm to replicate within the *Salmonella* Containing Vacuole (SCV) of host cells^9^. Because these genes function cooperatively, we refer to them collectively as the θ gene network. Importantly, each of these genes are highly conserved across *Salmonella enterica* serotypes indicating that they were acquired early in *Salmonella enterica’s* evolution^12^. Thus, we have proposed that the θ genes represent a “core” genetic network that allows *Salmonella* to replicate within animal cells.

Clearly, an organism engineered to express an artificially small gene repertoire should be less able to respond to environmental perturbations than its wildtype counterpart. Indeed, we found that although the θ genes were sufficient for intracellular replication of *S.*Tm *in vitro*, an engineered strain expressing just these five effector genes (called *S.*Tm θ) was avirulent in a mouse model of Typhoid-fever^9^. We then applied phenotype-guided genetic reassembly to determine that genes encoded by the *spv* operon, located on the *Salmonella* virulence plasmid, cooperate with the θ effector genes to enable *S.*Tm to survive and replicate in mice. The *spv* operon encodes three effector proteins, SpvB, SpvC, and SpvD. SpvB is an ADP-ribosyl transferase that suppresses host actin polymerization^13^. SpvC is a phosphothreonine lyase that inactivates Map kinases and JNK^14^. SpvD is a cysteine hydrolase that suppresses NF-kB mediated immune responses through an unknown mechanism^15^. Previous studies have shown that the *spv* effectors are associated with systemic infection and increased replication rates within macrophage *in vitro*^16,17^. However, unlike the θ effectors that are highly conserved across *Salmonella* species, the *spvBCD* effector genes have a restricted pattern of expression and are mostly found in non-typhoidal human serovars^12^. Thus, the evolutionary trajectory of some modern *Salmonella enterica* serovars, including *S.* Typhimurium (*S.*Tm), required the successful integration of *spv* effectors and their cooperation with the pre-existing θ effector gene network.

The identification of two independent gene networks (θ and *spv* operon) that were acquired at different time points of *Salmonella’s* evolution, yet cooperate to license systemic *S.*Tm infection of mice has raised several important questions about the integration of effector gene functions. For example, it is unclear why the θ gene network is insufficient for *S*.Tm pathogenesis in mice when these genes allow high levels of bacterial replication *in vitro*. What is the host bottleneck to infection by the *S.*Tm θ strain, and how did acquisition of the *spv* effectors overcome these defense mechanisms? It is also unclear what host cell types and tissues are primarily targeted by *S*.Tm expressing both the θ and *spv* operon effectors. Where do these two gene networks operate: in specialized immune cells or specific tissue environments? Finally, it is unclear if there are inflammatory mediators that suppress *S.*Tm θ replication. If so, how does expression of *spv* operon genes overcome these host immunological pathways? We set out to address these outstanding questions by combining the power of genome minimization with high content mass cytometry to track *S*.Tm replication within the host organism at single cell resolution.

In this study, we establish a comprehensive single cell atlas of *S*.Tm infection dynamics in a secondary lymphoid tissue. Our results reveal the spatiotemporal organization of bacterial transmission within a complex tissue and reveal how effector gene networks cooperate at the single cell level. We also uncover immunological bottlenecks targeted by the expression of the *spv* operon gene product SpvB that allows *S.*Tm to cause Typhoid-like disease in mice. These insights and the associated minimal strains, single cell tracking methodology, and atlas of *S.*Tm dissemination open doors to future characterization of effector gene network cooperation *in vivo* in the absence of confounding genetic and functional redundancies.

## Results

### Characterization of minimal *S.*Tm genetic strain phenotypes in cells and mice

We set out to establish an experimental framework to uncover the relationship between the θ and *spv* operon-encoded effector genes. As a benchmark for these studies, we first defined the growth parameters of wildtype *S.*Tm in a model epithelial cell line (HeLa), a model macrophage cell line (RAW 264.7), and disease-susceptible C57BL/6 mice. *S.*Tm replicated to high levels within LAMP-1 positive *Salmonella* Containing Vacuoles (SCVs) in HeLa cells and Raw 264.7 murine macrophage, similar to what has been observed in intestinal epithelial cells and primary macrophage, respectively^18,19^ (Fig. 1a and Extended Data Fig. 1a,b). In addition, C57BL/6 mice developed signs of Typhoid-like disease symptoms within two days of intragastric (i.g.) inoculation with *S.*Tm and severe signs of disease were apparent by 5 days post infection (dpi, Fig. 1f). These symptoms included reduced feeding behaviors, as quantified by monitoring food consumption after infection (Extended Data Fig. 1c). A consequence of the reduced food intake is lower activation of SREBP1 transcription factors in the liver (Extended Data Fig. 1d).

**Fig. 1:**
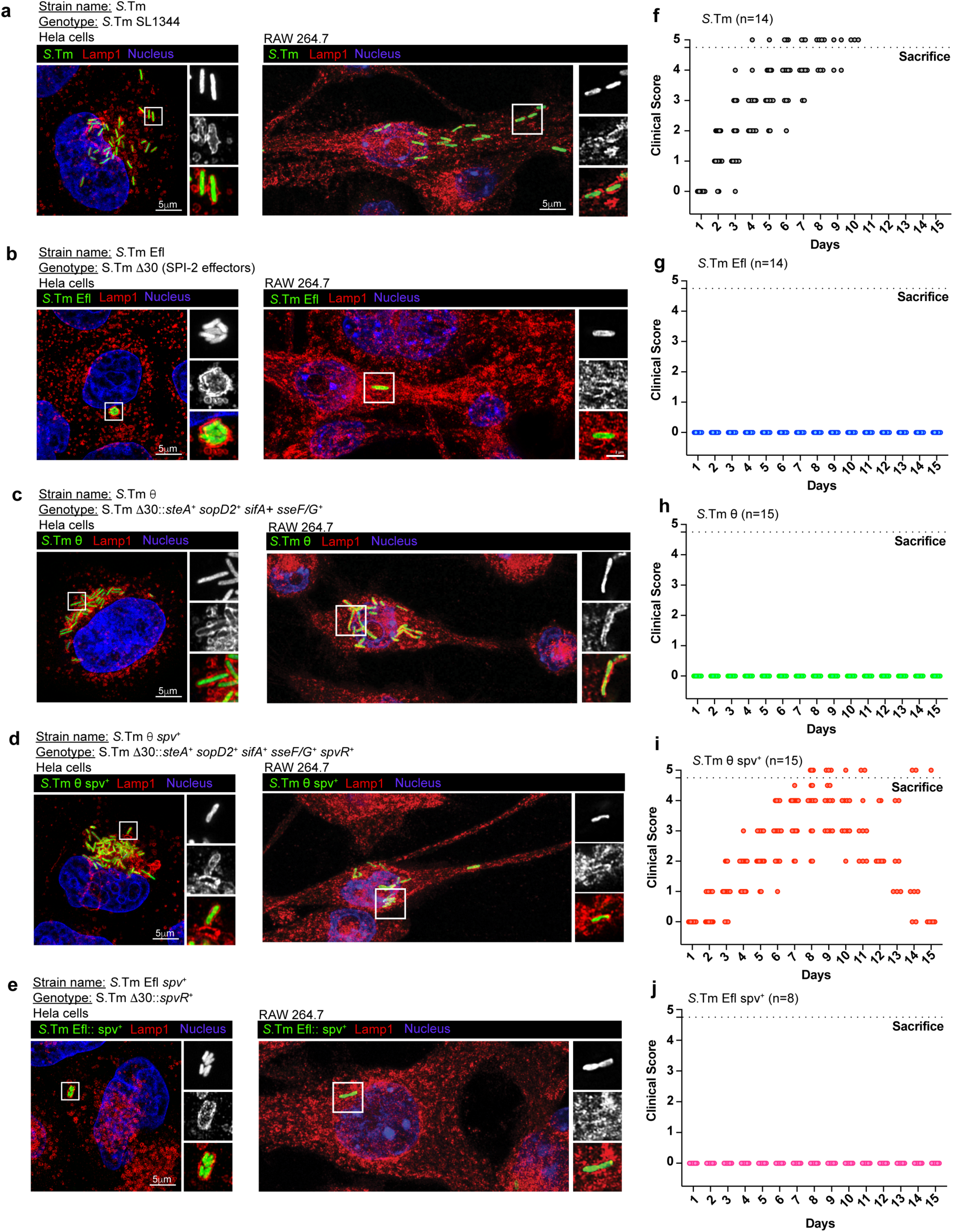
Establishing minimal strain phenotypes *in vitro* and *in vivo*. (a-e) Maximum intensity projection of confocal microscopy images of Hela cells (left panels) and Raw 264.7 murine macrophages (right panels) infected with GFP expressing bacterial strains: *S.*Tm (a), *S.*Tm Efl (b), *S.*Tm θ (c), *S.*Tm Efl *spv*^+^ (d) and *S.*Tm θ *spv*^+^ (e) (green) immunolabeled for LAMP1 (red) and DAPI (blue). Insets to the right are enlarged single slices from the white boxed region in the maximum intensity projection. The specific minimal effector genotypes of each strain is indicated above the images. (f-j) Clinical score of C57BL/6 mice infected i.g. with 1×10^9^ bacteria *S.*Tm (n=14) (f), *S.*Tm Efl (n=14) (g), *S.*Tm θ (n=15) (h), *S.*Tm θ *spv*^+^ (n=15) (i) and *S.*Tm Efl *spv*^+^ (n=10) (j). Experiments were conducted with three independent cohorts of mice (4-5 mice per experiment). Clinical score was measured on a scale of 0-5 where 0 represents no clinical symptoms and 5 represents severe clinical symptoms. Mice receiving a clinical score of 4.5 were sacrificed according to IACUC regulations.

We previously generated an *S.*Tm strain lacking all 30 SPI-2 effector genes (*S.*Tm Efl) that served as the second benchmark strain for comparative studies^9^. *S.*Tm Efl invaded tissue culture cells, likely because it harbors an intact SPI-1 T3SS that translocates effector proteins required for cell invasion^20^ (Fig. 1b). However, this strain was unable to replicate to high levels in either HeLa cells or Raw264.7 macrophage (Fig. 1b and Extended Data Fig. 1a,b). In addition, *S.*Tm Efl exhibited an abnormal SCV morphology in Hela cells wherein multiple bacteria were trapped in a single membrane compartment (Fig. 1b)^21^. The failure of *S.*Tm Efl to replicate intracellularly resulted in a severe virulence defect *in vivo* (Fig. 1g and Extended Data Fig. 1c,d) similar to what has been described for *S.*Tm strains lacking the SPI-2 T3SS apparatus^22^.

We previously identified five effector genes, *sopD2*, *steA*, *sifA*, *sseF, and sseG*, that are sufficient to support bacterial replication within SCVs in host cells^9^. This minimal strain (called *S.*Tm θ) displayed intracellular growth characteristics identical to wildtype *S.*Tm in Hela cells and Raw264.7 macrophage (Fig. 1c and Extended Data Fig. 1a,b). However, expression of these effectors alone were not sufficient for *S.*Tm to induce Typhoid-like disease symptoms or lethality in mice (Fig. 1h and Extended Data Fig. 1c,d). Thus, the gain-of-function phenotype of *S.*Tm θ compared to the S.Tm Efl benchmark strain can be attributed to the θ effector proteins driving SCV maturation and intracellular replication of the pathogen^9^. In contrast, the loss-of-function phenotype of *S.*Tm θ compared to the wildtype *S.*Tm benchmark strain in murine models of infection are likely due to the minimization of this pathogen resulting in loss of essential effector genes required for *Salmonella* to overcome physical or immunological bottlenecks imposed by the host organism.

Finally, we previously identified the *spv* operon effectors (*spvBCD*) as an independent gene network that cooperates with the θ effector genes^9^. To study the cooperation between these two gene networks, we generated the strain *S.*Tm θ *spv*^+^ (previously called STm Ω^9^) that expresses the θ effector genes along with the *spv* operon effector genes in the *S.*Tm Efl background. This strain has a functionally complete effector gene repertoire that, unlike *S.*Tm θ, is highly pathogenic in C57BL/6 mice (Fig. 1i and Extended Data Fig. 1a,b). Mice infected with *S.*Tm θ *spv^+^* develop Typhoid-like disease symptoms by 2 dpi with severe disease manifesting between 4-9 dpi (Fig. 1i). It should be noted that the onset of disease and the time to morbidity in *S.*Tm θ *spv^+^* infected mice was slightly slower than wildtype *S.*Tm (Fig. 1I). Nevertheless, both strains induced stereotypical sickness behaviors including anorexia and SREBP1 activation (Extended Data Fig. 1c,d). To ensure that *S*.Tm required co-expression of the θ genes and the *spv* operon, we tested the cellular and animal growth characteristics with *S.*Tm Efl *spv^+^*that only expresses the *spv* operon effector genes. Importantly, *S.*Tm Efl *spv^+^*does not replicate robustly in cell models and is avirulent in mice (Fig. 1e,j and Extended Data Fig. 1). Thus, θ and *spv* effector gene expression are necessary and sufficient for *S*.Tm virulence in mice.

### Mass cytometry detection of *S.*Tm within murine immune cells

Our *in vitro* studies suggest that the inability of *S*.Tm θ to colonize mice is not due to host restrictions on intracellular bacterial replication. Furthermore, addition of the *spv* operon effectors restores virulence to the *S*.Tm θ strain. To understand the complex relationship between these two effector gene networks *in vivo*, we set out to capture a system-wide view of *S.*Tm colonization at single cell resolution by mass cytometry (CyTOF). Briefly, an antibody recognizing *Salmonella* lipopolysaccharide (LPS) was conjugated with the lanthanide metal Dysoprosium-163 (Dy^163^) (Fig. 2a). This antibody recognized intracellular *S.*Tm as assessed by immunofluorescence microscopy and by conventional flow cytometry (Fig. 2b,c). Subsequently, we tested the metal conjugated antibody to detect *S.*Tm in primary immune cells. Mice were infected by intraperitoneal (i.p.) injection or intragastrically (i.g.) with lethal doses of *S.*Tm. Spleens were collected at 4 dpi and 7 dpi, respectively. Splenocytes were fixed and labelled with anti-CD45-Y^89^ antibody to detect the leukocyte common antigen. We then permeabilized and labelled these cells with LPS-Dy^163^. As expected, CD45^pos^ immune cells from uninfected mice exhibited low background levels of LPS-Dy^163^ signal (Fig. 2d). In contrast, high LPS signal (LPS^pos^) was detected in CD45^pos^ splenocytes from mice challenged with *S.*Tm by both the i.p. and i.g. routes of infection (Fig. 2d).

**Fig. 2:**
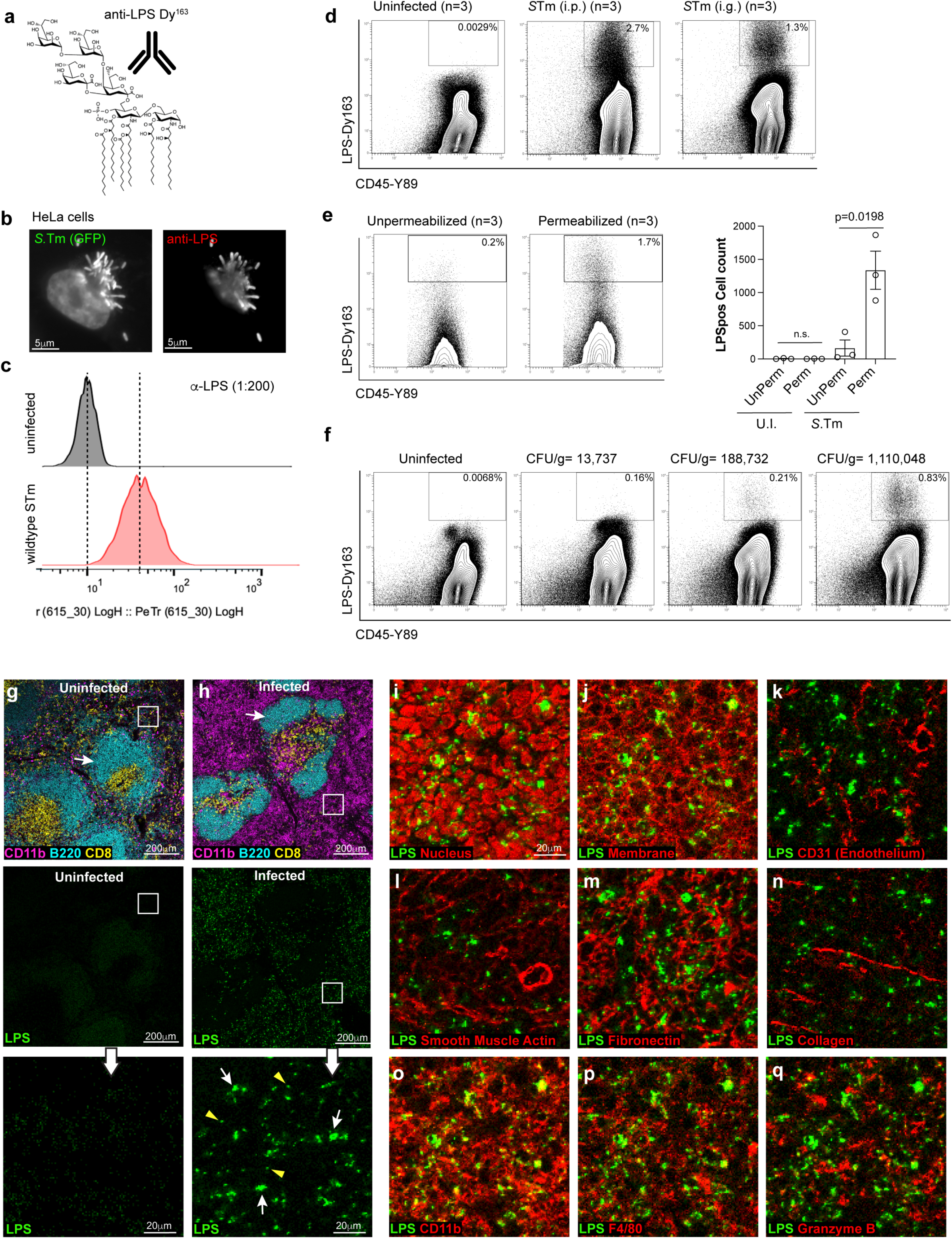
Development of mass cytometry for detecting *S.*Tm replication within murine immune cells. (a) Model of LPS and antibody conjugated to the Dy^163^ metal. (b) Immunofluorescence images of Hela cells infected for 18 hours with a GFP expressing *S.*Tm (MOI = 25) (left image) labeled by immunocytochemistry using the anti-LPS antibody (right image). (c) Flow cytometry histogram analysis of H1299 cells infected for 18 hours with *S.*Tm (MOI = 25) (green), labeled with the anti-LPS antibody and analyzed by flow cytometry. Uninfected cells (top), infected cells (bottom). (d) Flow plots showing mass cytometric analysis of murine splenocytes immunolabeled with anti- *S*.Tm LPS-Dy^163^ and anti-CD45-Y^89^ antibodies. C57BL/6 mice were uninfected (n=3) or infected intragastrically (i.g.) (1×10^9^ *S*.Tm bacteria) (n=3) or intraperitoneally (i.p.) (1×10^4^ *S*.Tm bacteria) (n=3) and sacrificed 7 dpi and 4 dpi, respectively. The percentage of cells within the LPS^pos^ region (box) is shown. (e) Flow plots from C57BL/6 mice infected with *S.*Tm as in (d). Mouse splenocytes were either permeabilized or unpermeablized with Perm-S buffer (see Materials and Methods) and then immunolabeled with anti-CD45-Y^89^ and anti-*S*.Tm LPS-Dy^163^ antibodies (left). Number of LPS^pos^ cells within total CD45^pos^ splenocytes collected from C57BL/6 mice from three independent experiments (right). Mean +/- SEM, Statistics: T-test. (f) Flow plots as in (d) of splenocytes collected from an uninfected mouse or from three individual i.g. infected mice. CFU/g of bacteria in the spleen of each mouse is shown above the plots. (n=1 mouse per CFU value, 1×10^9^ *S*.Tm bacteria/infection). (g-h) Imaging mass cytometry data collected on a Helios Mass Cytometer from spleen tissue sections of C57BL/6 mice uninfected (g) or infected i.g. (h) with 1×10^9^ *S*.Tm. Tissue sections are immunolabeled for CD11b-Sm^149^ (magenta), B220-Dy^162^ (cyan), and CD8-Yb^176^ (yellow) (top panels) LPS immunolabeling (green) in the same region as above (middle panels). Higher magnification of LPS immunolabeling in white boxed regions (lower panels). Arrows (white) show large colonies of *S*.Tm and arrowheads (yellow) show possible individual bacteria within cells. (i-q) Imaging mass cytometry data as in (g) showing anti-LPS signal (green) with the cell markers shown (red). DNA - Nuclei (i), plasma membrane (j), endothelial cells CD31 (k), smooth muscle actin (l), fibronectin (m), collagen (n), myeloid cells CD11b (o), myeloid cells F4/80 (p), and Cytotoxic lymphocytes Granzyme B (q).

An antibody directed at LPS could detect living bacteria and/or LPS particles that are ingested by phagocytes or displayed on the surface of antigen presenting cells^23^ (Extended Data Fig 2a). However, several lines of evidence indicate that the CyTOF methodology primarily detects living intracellular *S.*Tm. First, we did not detect LPS^pos^ signal in CD45^pos^ splenocytes isolated from mice infected with heat killed *S.*Tm by the i.p route of infection (Extended Data Fig. 2b). Thus, immune cells that ingest bacterial remnants do not constitute a significant proportion of the LPS^pos^ signal in *S.*Tm infected mice (Extended Data Fig. 2a,b). Second, detergent permeabilization was required to detect LPS^pos^ signal in infected CD45^pos^ splenocytes, indicating that CyTOF does not capture LPS particles displayed on the surface of antigen presenting cells (Fig. 2e). Finally, the total number of LPS^pos^ CD45^pos^ cells detected in spleen tissue increased linearly with the total number of living bacteria (CFU/g of spleen tissue) recovered from these animals (Fig.2f and Extended Data Fig 2c). Thus, the CyTOF methodology primarily detects living intracellular bacteria.

To further characterize the anti-LPS-Dy^163^ antibody, we applied imaging mass cytometry to spleen tissue samples from *S.*Tm infected mice displaying severe signs of Typhoid-like disease. Paraffin embedded tissue sections were stained with an antibody panel that included markers for canonical immune cell subsets (e.g. B-cells, T-cells, myeloid cells), markers for spleen architecture (e.g. endothelial cells, smooth muscle, and extra cellular matrix), and our anti-*S*.Tm LPS-Dy^163^. Hyperion mass cytometry revealed the splenic architecture at high resolution as denoted by the distinct zones of B-cells (B220^pos^) and CD8 T-cells (CD8^pos^) enriched within the white pulp of uninfected (Fig. 2g) and infected mice (Fig. 2h). Importantly, LPS^pos^ staining was apparent in tissues of mice infected with *S.*Tm (Fig. 2h below and inset), but not in uninfected samples (Fig. 2g below and inset). The LPS^pos^ signal was adjacent to the nucleus and within the plasma membrane borders of individual cells (Extended Data Fig. 3a,b) consistent with the perinuclear localization of intracellular *Salmonella* that has been reported *in vitro* (Fig. 2i,j)^24–26^. In addition, the LPS^pos^ signal in the cell ranged in size from what appeared to be potentially single bacteria (Fig 2h, inset, yellow arrowheads) to colonies likely consisting of multiple bacteria per cell (Fig. 2h, inset, white arrows) (Extended Data Fig. 3b). As previously observed, *S.*Tm was partially associated with splenic endothelial cells (Fig 2k), but *S.*Tm was not enriched in tissue regions dominated by smooth muscle cells, or structural components of the spleen and vasculature (Fig. 2l-2n)^27^. LPS^pos^ signal was also detected in tissue regions enriched with myeloid cells (CD11b^pos^ or F4/80^pos^), reflecting previous observations made with traditional immunohistochemistry^28^ (Fig. 2o,p). Together, these data support mass cytometry as a powerful tool to track the colonization dynamics of *S.*Tm at single cell resolution.

### A single cell atlas of *S.*Tm minimal strain infection of the spleen

We next used CyTOF to identify immune cell types infected by wildtype *S.*Tm and the three minimal strains (*S.*Tm Efl, *S.*Tm θ, and *S.*Tm θ *spv*^+^). We specifically analyzed bacterial colonization of the murine spleen since this organ houses diverse cell types from the hematopoietic lineage and since the SPI-2 T3SS is essential for spleen colonization by *Salmonella*. Mice were infected i.g. with each of the four *S.*Tm strains and spleens were collected at time points between 4 and 9 dpi. Isolated splenocytes were split into two samples. The first sample was used to measure the total intracellular bacterial burden within splenocytes of infected mice (CFU/g tissue). The second sample was labelled with a custom 26-plex antibody panel optimized for recognition of diverse innate and adaptive immune cell lineages and their activation states (see Material and Methods). Antibody labelled cells were then fixed, permeabilized, and stained with anti-LPS Dy^163^ to detect intracellular *S.*Tm. In total, we quantified marker expression on 17 million cells from five treatment groups: mock infection (n=3), wildtype *S.*Tm (n=11), *S.*Tm Efl (n=6), *S.*Tm θ (n=8) and *S.*Tm θ *spv*^+^ (n=6) using CyTOF (Extended Data Table 1). Quantification of the *S*.Tm CFUs/g of spleen tissue from each of the 34 mice is shown in Figure 3a.

**Fig. 3:**
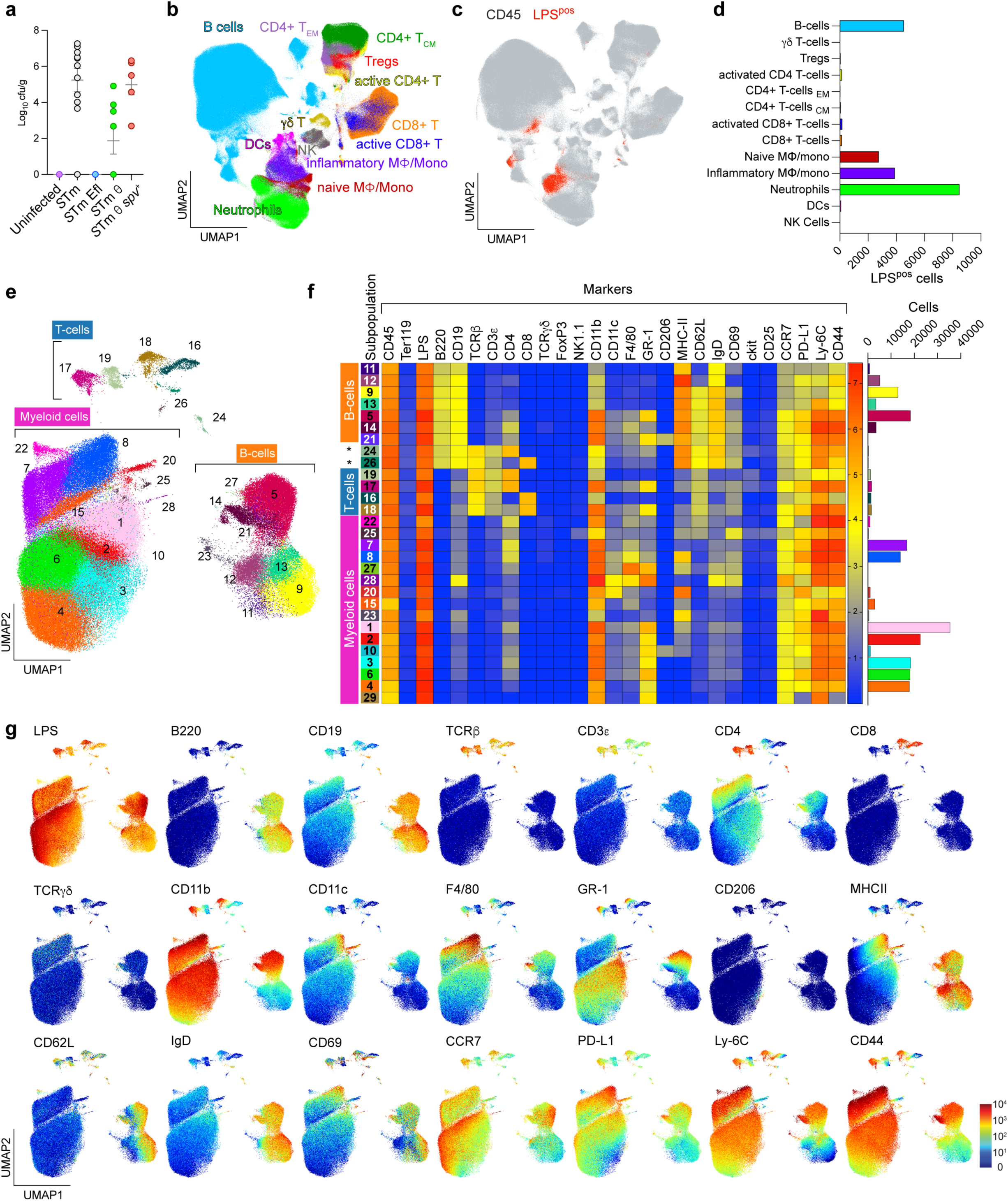
A single cell atlas of *S*.Tm infection. (a) Bacterial burden in spleens expressed as CFU/g for (n=34) C57BL/6 mice infected via intra gastric (i.g.) route with 1×10^9^ bacteria of the indicated *S.*Tm strains. Uninfected (n=3), *S*.Tm (n=11), *S*.Tm Efl (n=6), *S*.Tm θ (n=8), *S*.Tm θ *spv*^+^ (n=6). Mean +/- SEM. Individual mouse data is shown in Extended Data Table 1. (b) UMAP graph of 34 experimental mice in (a). Each dot in the UMAP graph represents an individual cell. Immune cell populations were color-coded and labelled based on manual gating (Extended Data Fig.4): B-cells (cyan), γδ T-cells (sage), Tregs (red), activated CD4+ T-cells (yellow), effector memory CD4 T-cells (lavender), central memory CD4+ T-cells (green), activated CD8+ T-cells (blue), CD8+ T-cells (orange), naïve macrophages/monocytes (rust), inflammatory macrophages/monocytes (purple), Neutrophils (neon green), Dendritic cells (pink), Natural Killer cells (gray). (c) UMAP graph of 34 experimental mice in (a) showing total CD45^pos^ cells (gray) and LPS^pos^ signal overlaid (red) for all 34 samples in the data set. (d) Total number of LPS^pos^ cells from (c) classified by immune cell types defined in (b). (e) UMAP visualization of the CD45^pos^ LPS^pos^ cells from all 34 experimental mice in (a) segmented by PhenoGraph into 29 subpopulations (sp). Colors and numbers correspond to each sp in the UMAP graph. (f) Heatmap showing normalized expression of the cell surface markers (top) in each of the PhenoGraph designated subpopulations. The total cell numbers in each subpopulation is shown (right). (g) UMAP graphs representing the CD45^pos^ LPS^pos^ total population from (e) where each graph shows the signal intensity for a single cell marker in individual cells.

Our first goal was to catalog all murine immune cell types infected with *S.*Tm regardless of the time of infection or the repertoire of effector proteins secreted by the pathogen. We evenly subsampled 35,714 CD45^pos^ cells from the spleen sample of each mouse (n=34) and used Uniform Manifold Approximation and Projection (UMAP) to reduce the dimensionality of the complex data set (Fig. 3b)^29^. For the analysis in Fig. 3b, broad immune cell phenotypes were assigned based on a manual gating strategy (Extended Data Fig. 4a). Viewing the distribution of LPS^pos^ signal (red) against the backdrop of CD45^pos^ immune cells (gray) revealed single cells infected by *S.*Tm on the UMAP graph (Fig. 3c). As expected, myeloid cells were the predominant infected cell types (Fig. 3d). However, we also noted populations of infected cells expressing B and T cell markers.

Because cells harboring intracellular bacteria are likely to exhibit non-canonical phenotypes, we next used a supervised algorithm (PhenoGraph) to more specifically classify the LPS^pos^ CD45^pos^ cells based on unique marker combinations^30^. We first gated on LPS^pos^ CD45^pos^ cells from the entire mouse cohort (Extended Data Fig. 4a, n=34) and then performed UMAP dimensionality reduction and PhenoGraph clustering (Fig. 3e). This approach revealed 29 *S.*Tm-infected subpopulations (sp) within the canonical myeloid cell, B-cell, and T-cell lineages (Fig. 3e and 3f). One of the largest groups of *S.*Tm infected cells of the myeloid lineage (CD11b^pos^ B220^neg^ TCRβ^neg^) were sp3, sp4, and sp6 (Fig. 3f). These subpopulations appeared to be functional substates of neutrophils based on their high expression of GR1, low expression of F4/80, and the absence of MHC class II. These data are consistent with neutrophils as a primary reservoir of *S.*Tm within the spleen^31^. In addition to neutrophils, *S*.Tm infected diverse macrophage and monocyte subpopulations. Among major *S*.Tm infected cell types (>10,000 cells detected), Sp8 had the highest expression level of the macrophage marker F4/80 (Fig. 3f). These cells also expressed high levels of MHC class II, revealing sp8 as the major antigen presenting macrophage cell type infected with *S.*Tm. The sp1 and sp2 populations also expressed high levels of MHC class II marker, but exhibited much lower levels of F4/80 compared to sp8. These subpopulations likely represent distinct substates of infected antigen presenting monocytes. Interestingly, sp10 was the only macrophage cell type to express CD206. Because CD206 is a marker of anti-inflammatory macrophage involved in tissue repair, the infected sp10 macrophage subset may be the cell type previously found to harbor drug-resistant *Salmonella* persisters^32^.

While neutrophils and macrophage dominated the infected myeloid cell population, there were smaller populations of cells that may have significant consequences for the progression of *S.*Tm infection. For example, sp20 exhibited the highest levels of both MHC class II and CD11c expression, indicating that these are dendritic cells infected with *S.*Tm. In addition, sp22 and sp25 were the only two myeloid cell types to express CD62L. CD62L, or L-selectin, is an adhesion molecule involved in mobilizing monocytes, B-cells, and T-cells across the endothelial cell barrier of blood vessels and has been previously implicated in *S*.Tm dissemination^33–37^. We therefore speculate that sp22 and sp25 are circulating blood monocytes that were infected by *S.*Tm shortly before taking residence in the spleen.

While myeloid cells had the highest burdens of *S.*Tm infection in general, LPS^pos^ signal was detected in 7 distinct subpopulations designated as B-cells (B220^pos^ TCRβ^neg^) (Fig. 3f). These B-cells could be further delineated into naïve (sp9, sp11, sp12, and sp13) or activated (sp5, sp14, and sp21) B-cells based on the absence (naïve) or presence (activated) of the markers CD11b^hi^ GR-1^pos^ F4/80^pos^. Individual B-cell subpopulations could be further distinguished from one another based on differential expression of CD4, CD206, MHC-II, CD62L, IgD, and CD69 (Fig. 3f,g). However, it is currently unclear if each subpopulation is a unique class of B-cells or are functionally related cells at different stages of activation.

Finally, CD4^pos^ and CD8^pos^ T-cells (sp16-sp19) infected with *S.*Tm were detected by CyTOF (Fig. 3e). It should be noted however that T-cells represented less than 3% of the total LPS^pos^ CD45^pos^ population suggesting that this is a minor niche of *S.*Tm colonization (Fig. 3d,f). We did not find evidence of infected γδ T-cells (TCRγδ^pos^) or regulatory T-cells (FoxP3^pos^ CD25^pos^) suggesting that the LPS^pos^ signal within CD4^pos^ and CD8^pos^ T-cells represents bonafide intracellular *S.*Tm. It was difficult to classify these infected T-cells into effector or memory cells since all cells infected expressed varying levels of the CD69 and MHC-II markers that are typically used to differentiate these cell phenotypes. However, we could distinguish two distinct classes of infected CD4^pos^ and CD8^pos^ T-cells based on differential marker expression. For example, sp17 and sp18 expressed CD4 and CD8 markers along with high levels of CD11b and GR-1 (Fig. 3f,g), whereas levels of these markers were not elevated in sp16 and sp19. Also, sp17 and sp18 exhibited higher LPS^pos^ signal suggesting that individual cells in these subpopulations have higher burdens of *S.*Tm compared to their sp16 and sp19 T-cell counterparts.

### Single cell infection dynamics of *S.*Tm colonization of the spleen

After generating a comprehensive map of bacterial infection from mice infected with multiple strains of *S.*Tm (Fig. 3e) we then directly compared the landscape of PhenoGraph-identified immune cell subpopulations (sp) from individual mice infected with wildtype *S.*Tm to the landscape of immune cell subpopulations infected with the two *S.*Tm minimal strains (Fig. 4). Importantly, our data set included CD45^pos^ LPS^pos^ cells isolated from mice with varying levels of spleen infection based on bacterial CFU/g isolated from spleens (Extended Table 1). We reasoned that mice with low CFUs in the spleen were sacrificed at an earlier stage of disease progression compared to mice with higher CFUs (Fig. 2f, Extended Data Fig. 2c). Thus, we were able to construct a time-resolved depiction of immune cell populations infected with wildtype *S*.Tm at early to late stages of spleen colonization. We then used this depiction as a benchmark to understand the contributions of the θ and *spv* effector gene networks to bacterial spleen colonization (Fig. 4a-c).

**Fig. 4:**
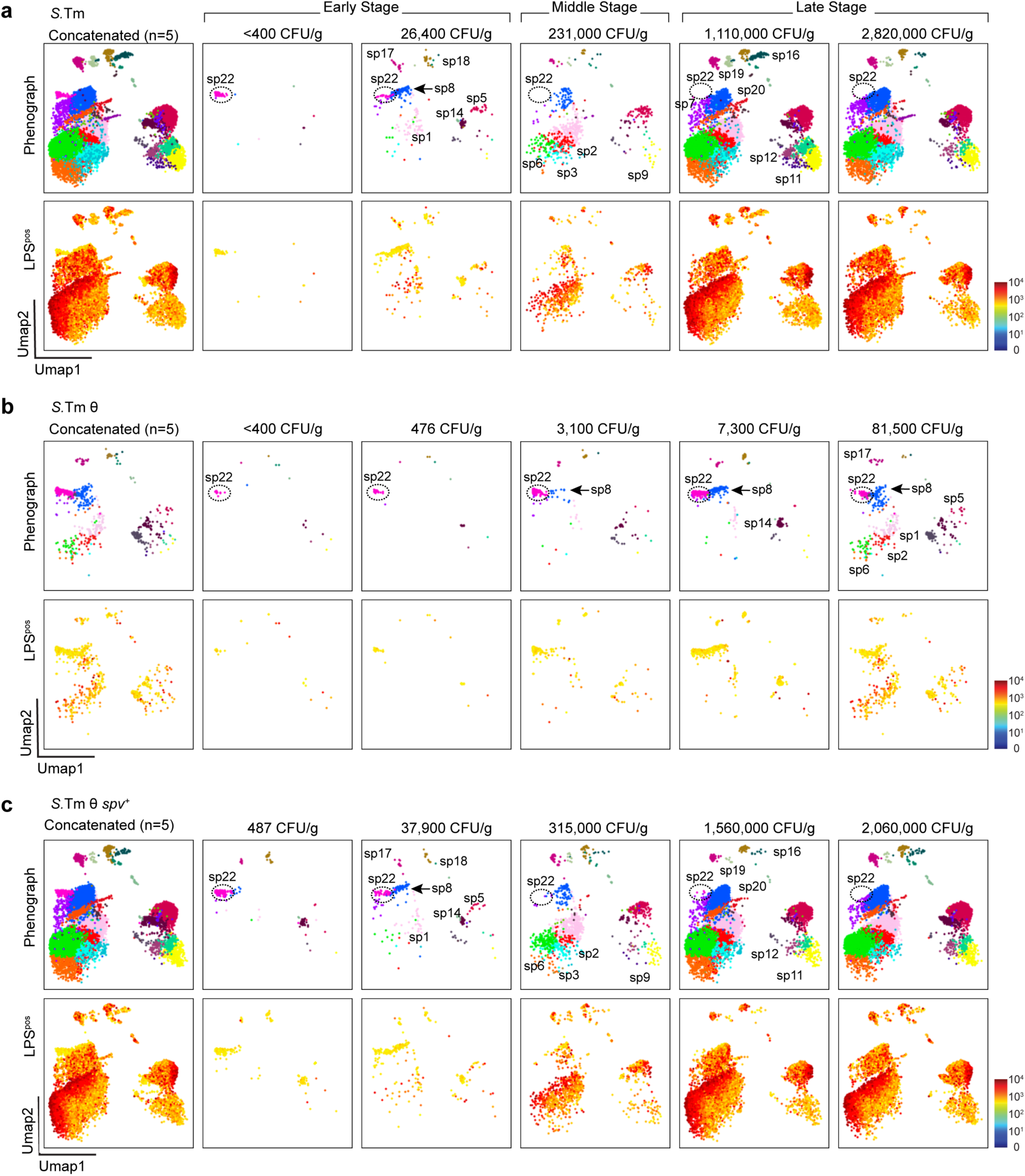
Spatiotemporal analysis of SPI-2 effector gene cooperation. (a-c) UMAP visualization of CD45^pos^ LPS^pos^ cells from individual mice infected i.g. (1×10^9^ bacteria) with S.Tm (a), S.Tm θ (b), and S.Tm θ *spv^+^*(c) as described in Fig. 3e. The left plots show the UMAP of 5 concatenated samples (n=5 mice). The right plots show the UMAPs for each of the 5 individual mice ordered from lowest (left) to highest (right) bacterial burden in the spleen (CFU/g). The top UMAP graph (in a,b and c) shows PhenoGraph sub-populations labelled and colored as in Fig. 3e,f. The bottom UMAP is a heatmap of LPS-163^Dy^ labeling intensity.

We first analyzed the immune cell subpopulations from five mice infected with wildtype *S*.Tm that allowed us to define early, middle, and late stages of bacterial colonization (Fig. 4a). We found that sp22 was the predominant LPS^pos^ cell type observed at the earliest stage of *S.*Tm infection (<400 CFUs/g tissue). As described above, sp22 cells are CD62L^pos^ circulating monocytes that are likely infected by *S*.Tm within the bloodstream prior to occupying the spleen (Fig. 3f and Fig. 4a). As *S.*Tm burden increased from <400 CFU/g to 26,400 CFU/g, we observed LPS^pos^ signal within the sp8 antigen-presenting macrophage population (blue; CD11b^pos^ F4/80^pos^ MHCII^pos^) as well as some B- and T-cells. The sp8 macrophage were notable because they did not express CD62L indicating that appearance of these cells occurred after sp22 arrival in the spleen. Interestingly, as *S.*Tm infection progressed to 231,000 CFU/g (middle stage), sp22 cells were no longer detected (Fig. 4a). These monocytes were never detected again even at the highest levels of *S.*Tm in the spleen. The sp22 collapse correlated with expansion of *S*.Tm in activated macrophage (sp8), antigen presenting monocyte and macrophage (sp1 and sp2), neutrophils (sp3 and sp6) and naïve B-cells (sp5, and sp9). Importantly, we did not observe further *S*.Tm expansion into either T-cells or B-cells between the early and middle stages of infection. Thus, the emergence of new LPS^pos^ signal in the myeloid cell lineage is likely due to cell-to-cell dissemination of the pathogen within physically restricted regions of the spleen. Finally, as wildtype *S.*Tm burdens surpassed 1×10^6^ CFU/g (late stage), the LPS^pos^ cellular landscape reached a steady state distribution accounting for infection of 28 out of the 29 PhenoGraph-defined subpopulations (Fig. 4a).

### Cooperation between θ and *spv* operon effector genes drive *Salmonella* cell-to-cell dissemination

The cellular progression of *S.*Tm through distinct sp’s of host cells provided opportunity to examine the function of the θ and *spv* effector gene networks and their mechanisms of cooperation in a single cell time-resolved system. When analyzing LPS^pos^ signal from individual mice we noted that most *S.*Tm θ infections failed to progress past an early stage colonization pattern (Fig. 4b). CD62L^pos^ circulating monocytes (sp22 cells) were the major infected cell type even in mice with bacterial colonization below the limit of detection by tissue sampling. We also noted that the infected sp22 cell population did not collapse at any point during *S.*Tm θ infection (Fig. 4b). These data indicate that replication competent *S.*Tm can infect and survive within CD62L^pos^ circulating monocytes for long periods of time. However, without the cooperation of other effector proteins, namely the *spv* operon, *S.*Tm θ is unable to efficiently spread from this cell type and/or survive within tissue resident immune cells. Thus, it appears that sp22 cells create a bottleneck through which *S*.Tm infection does not efficiently pass without the *spv* effectors. While this bottleneck at early stages of *S*.Tm θ infection was predominantly sp22-mediated, at the highest CFU/g achieved by this strain (81,500 CFU/g), there was evidence of infection of macrophage (sp8 cells) as well as neutrophils (sp1, sp2, sp6). However, the level of LPS^pos^ signal within these cell populations were quite low compared to that achieved by wildtype *S.*Tm, suggesting that these cells actively suppress replication of *S.*Tm θ.

Remarkably, *S.*Tm θ *spv*^+^ exhibited cell colonization dynamics nearly identical to wildtype *S.*Tm (Fig. 4c). Not only did the initially infected CD62L^pos^ circulating monocyte (sp22) population collapse at a similar stage of infection, but *S.*Tm θ *spv*^+^ replicated to high levels in activated macrophage, monocytes and neutrophils. Thus, the combination of θ effector genes and the *spv* operon effectors are the primary driving mechanism of *S.*Tm colonization of the spleen. These effectors allow replication competent bacteria to overcome host-imposed bottlenecks that block either cellular egress from circulating monocytes (sp22) or survival in other splenic cell types. The *spv* operon also directs expansion of the pathogen into multiple cellular niches through a consistent trajectory of dissemination (Fig. 4).

### Host immune cell response to *Salmonella* minimal strains

Our single cell data not only captured the trajectory of *S*.Tm cell-to-cell dissemination within the spleen, but also the tissue immune response to each *S.*Tm strain expressing different effector gene combinations. To directly compare the host response to infection between the minimal strains (*S.*Tm, *S.*Tm Efl, *S.*Tm θ and *S.*Tm θ *spv^+^*), we concatenated the total CD45^pos^ cell population from three individual mice infected with each of these *S.*Tm strains at 7 dpi (Extended Data Table 1). UMAP visualization of these data allowed us to evaluate changes in immune cell population marker expression in mice infected for a common duration (Fig. 5a). We also quantified the total number of immune cell subsets in the spleen by back calculating from the percentage of live cells in the collected sample of total CD45^pos^ cell population (Fig. 5b). In general, we observed either an increase in cell number, a decrease in cell number or an alteration in activation state for nearly every immune cell type in response to wildtype *S*.Tm infection (Fig. 5a,b). For example, *S.*Tm induced an influx of neutrophils, macrophage, and active CD8^pos^ T-cells into splenic tissue (Fig. 5b). We also observed a retraction of the effector memory and central memory T-cell populations under these conditions. While there was no change in total B-cell numbers within the spleen, these cells were highly activated by *S.*Tm infection (Fig. 5a,b). Interestingly, the UMAPs from mice infected with *S*.Tm Efl or *S*.Tm θ strains show an overall immune cell pattern that is similar to uninfected mice (Fig. 5a). This is consistent with the low levels of spleen colonization (Extended Data Table 1) and the low clinical scores (lack of symptoms) caused by infection with these strains (Fig. 1). In contrast, the host immune cell response to *S.*Tm θ *spv^+^* infection was nearly identical to wildtype *S.*Tm. These data reinforce our conclusion that co-expression of the θ and *spv* operon effectors is the primary driver of cell-to-cell dissemination of *S.*Tm, which consequently drives the host immune cell responses to infection in the spleen.

**Fig. 5:**
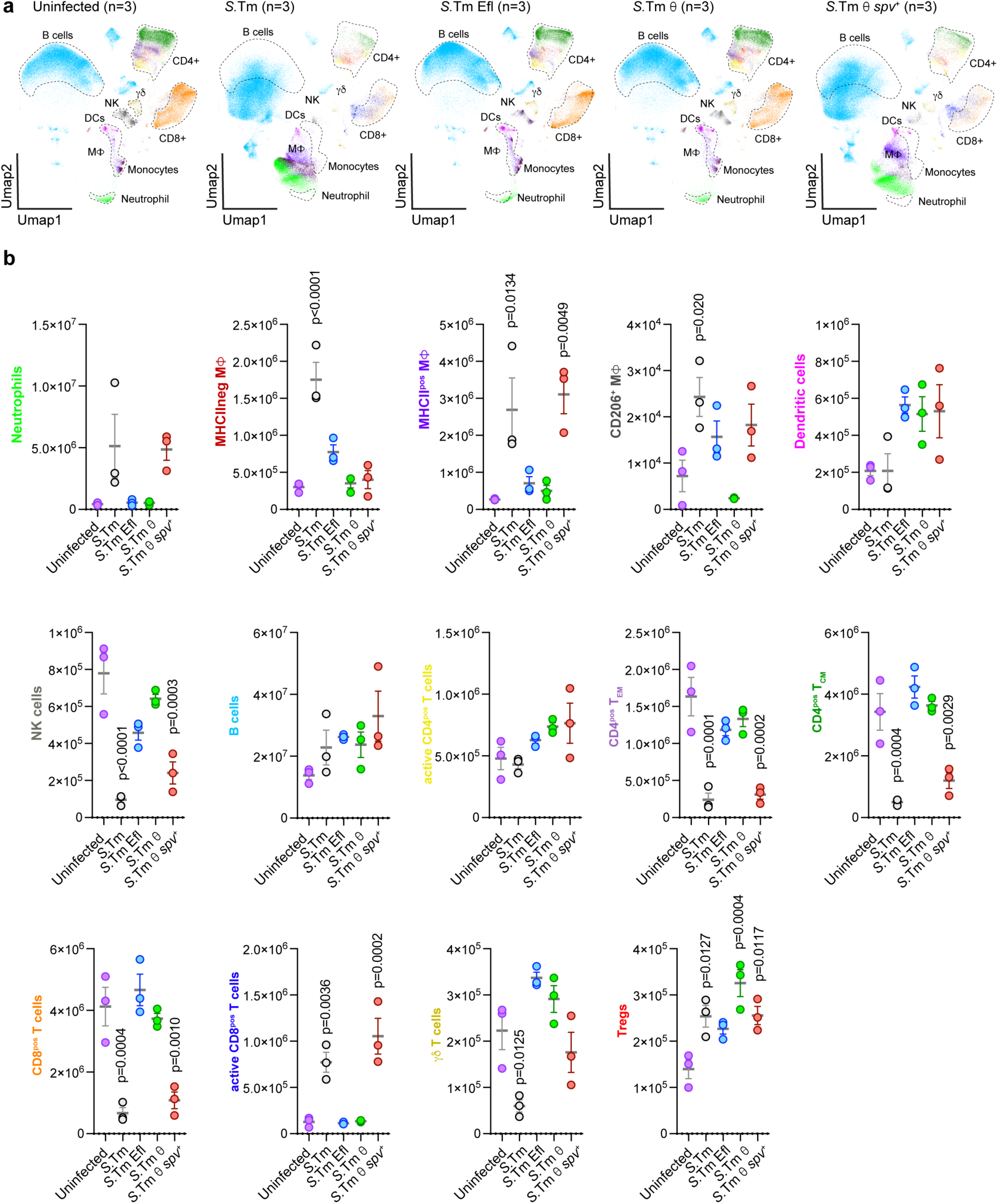
Quantitative analysis of immune cell subsets during *Salmonella* infection. (a) UMAP graphs of the entire splenocyte population were used to analyze the host immune response to different *S*.Tm strains. 3 uninfected mice, as well as 3 mice infected i.g. with 1×10^9^ bacteria with each of the strains *S.*Tm, *S.*Tm Efl, *S.*Tm θ, or S.Tm θ *spv*^+^ and sacrificed at 7 dpi were included in this analysis (see Extended Data Table 1). Each UMAP is a concatenation of the 3 mice infected with the indicated *S.*Tm strain. Dotted lines represent the position in the UMAP graph of cell populations in uninfected murine samples. (b) Total number of cells in spleen samples for each immune cell type shown in (a). Mean +/- SEM (n=3 mice/group). Statistics: one-way ANOVA with Dunnett’s multiple comparison. p<0.05 compared to uninfected control is shown.

### Genetic conflict between *S.*Tm effectors and the Myd88 and IFN-γ signaling pathways

So far our single cell data indicates that evolutionary pairing of the θ effector genes with the *spv* operon allowed *Salmonella* to survive and disseminate within the immunological environment of mice. While the θ genes are clearly involved in modulating host factors required for *S*.Tm replication in the vacuole environment of immune cells (Fig. 1)^9^, the role of the *spv* operon effectors in promoting tissue colonization and Tyhpoid-like disease in mice is less clear. We reasoned that the *spv* effectors might protect the pathogen from a major branch of the immune system. Because *S.*Tm θ is avirulent in mice whereas *S.*Tm θ *spv^+^* is highly pathogenic, it is possible that elimination of host genes that exert pressures on *Salmonella* to express the *spv* effector genes could rescue the virulence defects of the attenuated *S.*Tm θ strain (Fig. 6). To test this, we infected mice lacking genes known to be critical for controlling the host immune response to *S.*Tm infection. For example, *S.*Tm is recognized by Toll-like receptors (TLRs), which activate NF-kB transcriptional responses through the Myd88 signaling pathway^38,39^. As expected, *S.*Tm was highly pathogenic in mice lacking Myd88 (*Myd88^-/-^*) (Fig. 6a). In contrast, the *S.*Tm Efl strain lacking all 30 SPI-2 effector genes had no effect on *Myd88^-/-^*mice indicating that *Myd88^-/-^* mice are not simply susceptible to an attenuated *S.*Tm genetic variant (Fig. 6b). Remarkably, while wildtype C57BL/6 mice were completely resistant to *S.*Tm θ infection, *Myd88^-/-^* mice were infected by this minimal strain (Fig. 6c). To determine if the virulence of *S.*Tm θ in *Myd88^-/-^* mice was simply caused by general immunosuppression, we also infected mice lacking *Nlrc4* with *S.*Tm strains. NLRC4 is a key signaling adaptor for the NLR-family apoptosis inhibitory proteins (NAIPs) that detect the *Salmonella* T3SS and flagellin^40–42^. As expected, *S.*Tm was highly pathogenic to *Nlrc4^-/-^* mice whereas *S.*Tm Efl was avirulent (Fig. 6a,b). Unlike in *Myd88^-/-^* mice, however, *S.*Tm θ had no effect on *Nlrc4^-/-^* mice (Fig. 6c).

**Fig. 6:**
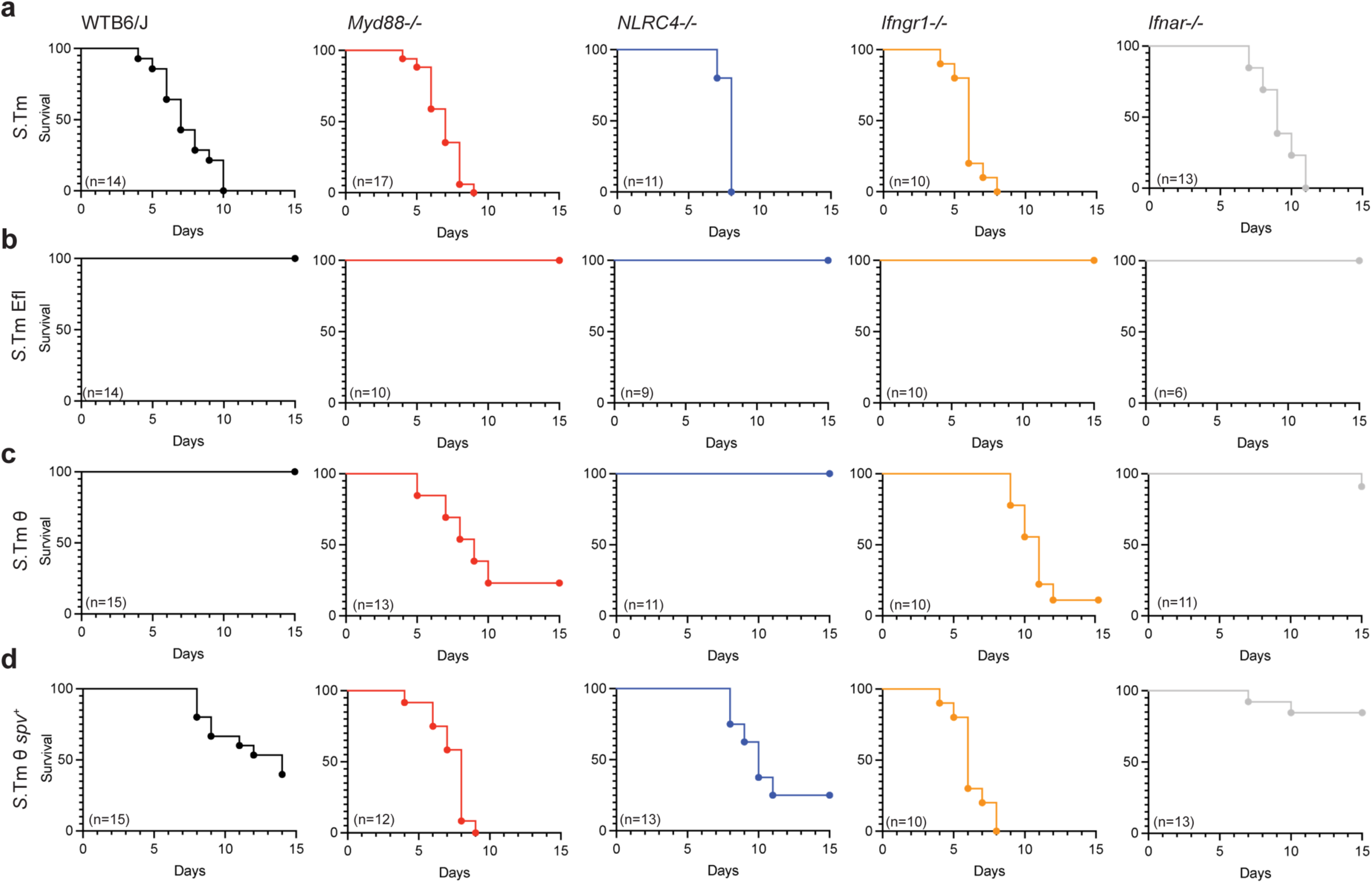
*Salmonella* minimal effector network contributes to pathogenesis in mouse knock out infection models. (a-d) Kaplan-Meier survival curves for the indicated mouse genetic strains (from left to right) C57BL/6 WT (black), *Myd88*^-/-^ (red), *Nlrc4*-/- (blue), *Ifngr1*^-/-^ (yellow), *Ifnar*^-/-^ (gray) infected i.g. with a dose of 1×10^9^ of the indicated strains: wild-type *S.*Tm (a), *S.*Tm Efl (b), *S.*Tm θ (c), *S.*Tm θ *spv*^+^ (d). The total number of mice from two experimental cohorts are indicated on the bottom left of all graphs. Mice were monitored for survival once daily and sacrificed accordingly to UTSW IACUC guidelines.

We next tested if cooperation between the θ and *spv* operon effectors is necessary for *S*.Tm to overcome a second major branch of the host immune system, the interferon response. Mice lacking signaling through the Type I (IFNα/β, *Ifnar1*^-/-^) or Type II IFN (IFNγ, *Ifngr1*^-/-^) receptors were infected with each of the *S.*Tm strains. As expected, *S.*Tm Efl was avirulent in both *Ifnar1*^-/-^ or *Ifngr1*^-/-^ mice (Fig. 6a,b). In contrast, *S.*Tm θ was highly pathogenic in *Ifngr1*^-/-^ mice, similar to what was observed in *Myd88*^-/-^ mice. Interestingly, *Ifnar1*^-/-^ mice were resistant to *S.*Tm θ induced disease, but were also resistant to *S.*Tm θ *spv*^+^ revealing a more complex relationship between *Salmonella* effector genes and the Type I IFN signaling pathway ^43,44^. Nevertheless, these data together indicate that co-expression of the θ and *spv* effectors are required for intracellular *S.*Tm to overcome the immunological bottlenecks imposed by murine TLR and IFN-γ signaling pathways.

### The θ and *spv* effectors promote *S*.Tm colonization of lymphoid tissues protected by the TLR-signaling pathway

The TLR/Myd88 and IFNγ pathways protect multiple tissues from bacterial infection^45^. We therefore sought to identify specific locations of conflict between the *spv* effector genes and the TLR/Myd88 and IFNγ pathways. Mice were infected i.g. (1×10^9^ CFU) with each *S*.Tm minimal strain and bacterial colonization levels in individual tissues were examined 4 dpi. Previous studies have shown that shortly after colonizing the small intestine, wild type *S.*Tm invades follicle-associated epithelial cells (e.g. M cells) which act as an interface between the intestinal lumen and the underlying sub-epithelial dome of the Peyer Patches (PPs)^46,47^. *S.*Tm and each of the minimal strains replicated to high levels within the luminal compartment of the caecum, confirming that the SPI-2 effector genes do not play a significant role in microbial competition within the gut (Fig. 7a)^48^. Interestingly, 80% of mice exhibited PP colonization by wildtype *S.*Tm whereas 20% of mice successfully defended this niche, despite high pathogen burdens in the lumen. In contrast, we did not observe *S.*Tm Efl colonization of PPs supporting a critical role of SPI-2 T3SS effectors in colonizing PPs (Fig. 7a)^22^. Similarly, we found that *S.*Tm θ was unable to colonize PPs whereas *S.*Tm θ *spv*^+^ exhibited a biphasic colonization pattern, similar to wildtype *S.*Tm. Thus, cooperation between these two effector gene networks is required for local tissue penetration of the gut mucosa.

**Fig. 7:**
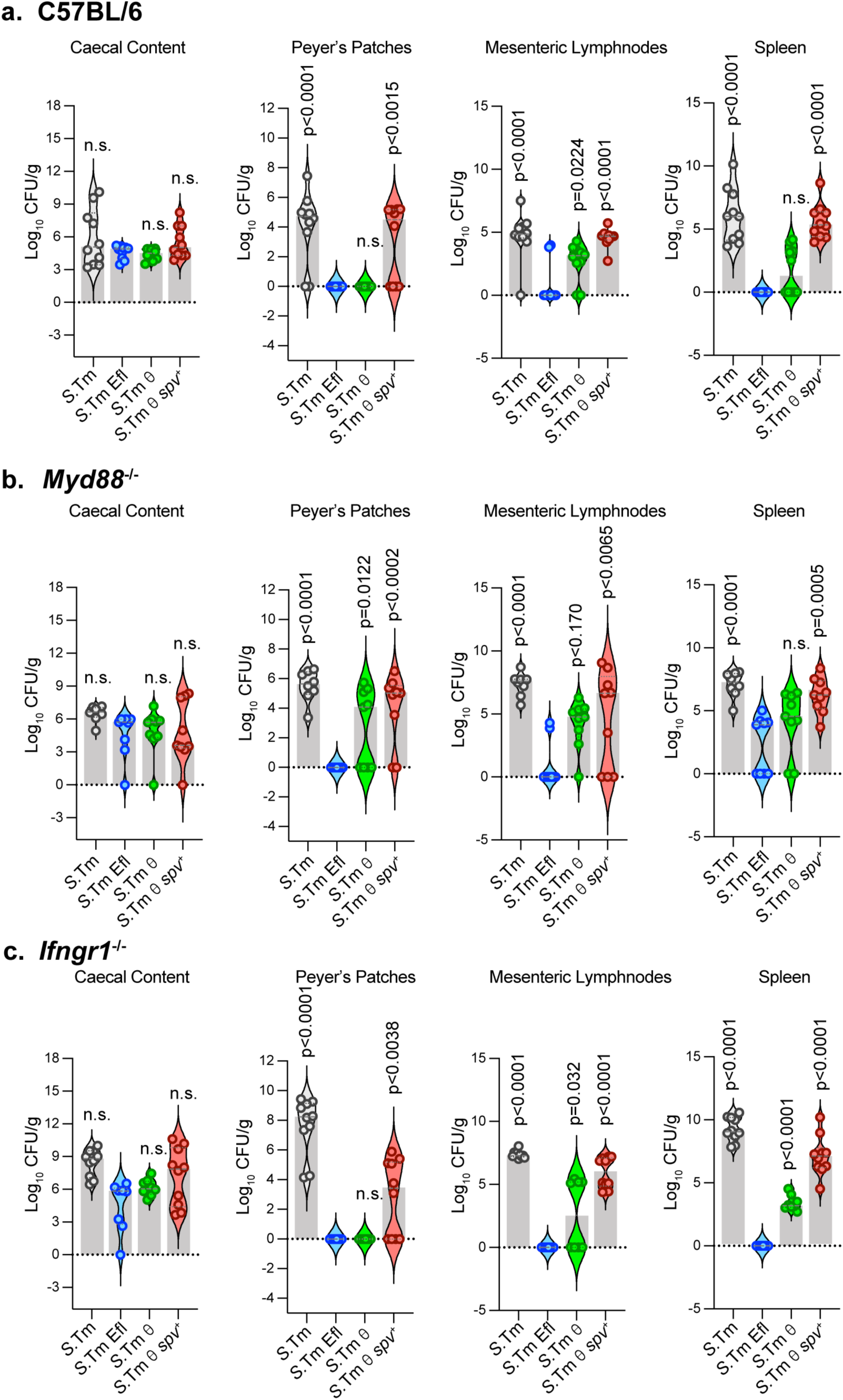
Bacterial burden of *Salmonella* minimal effector network strains in tissues during infection. (a-c) Bacterial burden (CFU/g) in caecal content, Peyer’s Patches, mesenteric lymph nodes and spleen from mice infected i.g. with 1×10^9^ bacteria of the indicated strains (*S.*Tm, *S.*Tm Efl, *S.*Tm θ, *S.*Tm θ *spv*^+^) and sacrificed at 4 dpi. C57BL/6 WT (a) C57BL/6 WT mice infected with strains: *S.*Tm (n=10), *S.*Tm Efl (n=10), *S.*Tm θ (n=10), and *S.*Tm θ *spv*^+^ (n=10). (b) *Myd88*^-/-^ mice infected with strains: *S.*Tm (n=8), *S.*Tm Efl (n=9), *S.*Tm θ (n=9), and *S.*Tm θ *spv*^+^ (n=9). (c) *Ifngr1*^-/-^ mice infected with strains: *S.*Tm (n=10), *S.*Tm Efl (n=8), *S.*Tm θ (n=8), and *S.*Tm θ *spv*^+^ (n=10). Mean (gray bars) +/- SEM. Statistics: one-way ANOVA with Dunnett’s multiple comparison. p<0.05 as compared to *S*.Tm Efl control are shown.

*S.*Tm can also cross the epithelial cell barrier into the Lamina Propria (LP) through dendritic cell (DC) sampling ^49,50^. Once the sub-mucosa is breached, *S.*Tm is transported by DCs through the lymphatic system to the mesenteric lymph nodes (MLNs) and from here the pathogen disseminates to systemic tissues (e.g. spleen). Both *S.*Tm Efl and *S*.Tm θ exhibited lower colonization rates of the MLNs and spleen compared to wildtype *S.*Tm and the *S.*Tm θ *spv^+^* strain (Fig. 7a). Thus, the combination of θ and *spv* effector genes is required for bacterial growth and survival within each lymphoid tissue tested.

We next sought to determine if the θ and spv operon effectors counteract the TLR or IFNγ signaling pathways at specific tissues along the route of *S.*Tm infection. *Myd88^-/-^* and *Ifngr1^-/-^*mice were infected with each of the *S.*Tm strains and lymphoid tissues were harvested at 4 dpi. As we observed in wildtype mice, *S.*Tm and *S.*Tm θ *spv^+^*strains colonized the gut lumen, PPs, MLNs, and spleen to high levels in *Myd88^-/-^*and *Ifngr1^-/-^* mice (Fig. 7b,c). However, examination of *S.*Tm θ colonization under these conditions was revealing. The most striking finding was that unlike wildtype mice that protect the PPs from *S.*Tm θ, *Myd88^-/-^* mice failed to inhibit PP infection, allowing *S.*Tm θ to colonize the PPs to high levels (Fig. 7a,b). Interestingly, *Ifngr1^-/-^* mice protected the PP niche from *S*.Tm infection, indicating that the TLR signaling pathway is the primary target of the *spv* operon effectors in this lymphoid tissue environment (Fig. 7c). Although we noted some interesting differences in the ability of *S.*Tm θ to infect MLNs and spleen in *Myd88^-/-^*and *Ifngr1^-/-^* mice, infection levels remained lower than wildtype *S*.Tm and further studies are needed to define the function of the *spv* operon at these distal sites of infection (Fig. 7b,c). Nevertheless, these studies support our conclusion that combined expression of the θ genes and *spv* operon are sufficient for *S*.Tm dissemination within lymphoid tissues that are protected by TLR-mediated signaling pathways.

### The *spvB* effector overcomes tissue specific bottlenecks to infection

Finally, we sought to determine if there is a specific effector in the *spv* operon that can overcome host immunological bottlenecks to promote *S.*Tm colonization of lymphoid tissues. We generated mutant *S.*Tm θ *spv+* strains with single deletions of *spvA*, *spvB*, *spvC*, and *spvD*. Remarkably, *S.*Tm θ *spv+* lacking *spvA*, *spvC*, or *spvD* was highly virulent, whereas *S*Tm θ *spv+* Δ*spvB* showed attenuated colonization in the Peyer’s Patches, liver, and spleen (Fig. 8a). Importantly, complementation of *S*Tm θ *spv+* Δ*spvB* by expression *spvB* from the PhoN locus (*S.*Tm θ spv^+^Δ*spvB phoN::spvB*) restored the virulence of this strain, indicating that knocking out *spvB* did not introduce polar effects on the co-transcribed *spv* operon (Fig. 8a). SpvB functions as an ADP-ribosyl transferase that post-translationally modifies host cellular G-actin^13,51^. We therefore conclude that dissemination of *S.*Tm between different cell types during the course of an *in vivo* infection requires θ effector protein regulation of SCV membrane dynamics combined with SpvB*-* mediated suppression of host actin polymerization.

**Fig. 8:**
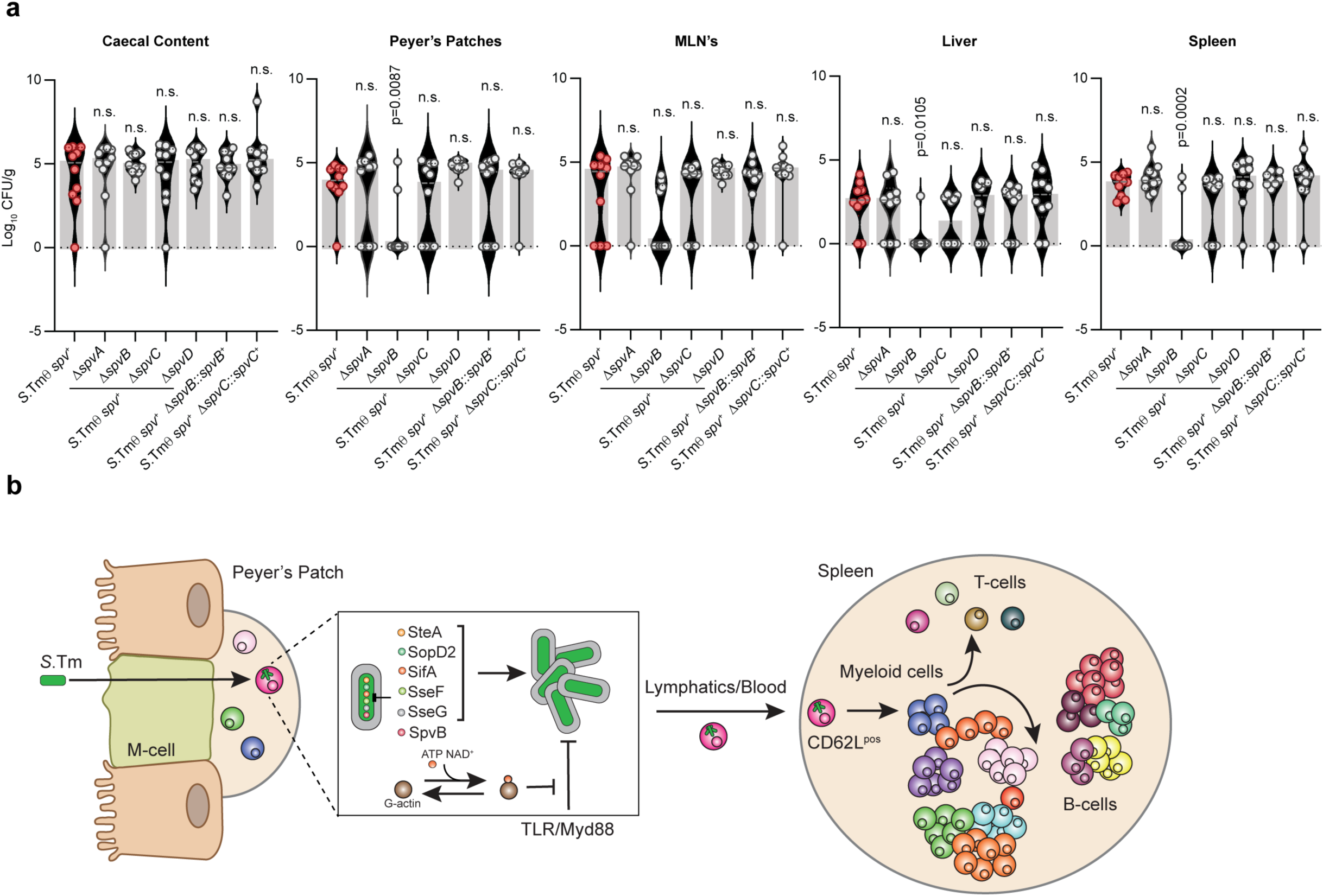
The *spv* operon effector *SpvB* contributes to *S*.Tm dissemination in tissues. (a) Bacterial burden (CFU/g) in caecal content, Peyer’s Patches, mesenteric lymph nodes, liver, and spleen from C57BL/6 mice infected via the intra gastric (i.g.) route with 1×10^9^ of the indicated strains and sacrificed at 4 dpi. *S.*Tm strains: *S*.Tm θ *spv*^+^ (n=10), *S*.Tm θ *spv*+Δs*pvA* (n=10), *S*.Tm θ *spv*^+^Δs*pvB* (n=10), *S*.Tm θ *spv*^+^Δs*pvC* (n=10), *S*.Tm θ *spv*^+^Δs*pvD* (n=10), *S*.Tm θ *spv*^+^Δs*pvB*::*spvB*^+^ (n=10), *S*.Tm θ *spv*^+^Δs*pvC*::*spvC^+^* (n=10). Mean +/- SEM. Statistics: one-way ANOVA with Dunnett’s multiple comparison. p<0.05 compared to *S*.Tm θ *spv*^+^ control are shown. (b) Model of *S*.Tm colonization dynamics and the role of effector gene networks in systemic infection. *S.*Tm colonizes the gastrointestinal tract and penetrates the intestine through M-cells resulting in colonization of gut associated lymphoid tissues including Peyer’s Patches. The θ effector proteins SteA, SopD2, SifA, SseF, and SseG promote intracellular replication and SCV division of *S*.Tm within immune cells whereas SpvB ADP-ribosylates G-actin, which is responsible for counteracting Myd88-mediates suppression of infection. *S*.Tm is likely transported to the spleen by CD62L^pos^ monocytes, where this cell type releases the pathogen or differentiates into macrophage within the murine spleen. The combination of θ and *spv* effectors are required for cell-to-cell spread through a common trajectory of dissemination.

## Discussion

In this study, we have coupled a genome minimization strategy with single-cell mass cytometry to uncover host-pathogen conflicts that are hidden beneath layers of genetic redundancy (Fig 8b). Specifically, we adapted Cytometry by Time of Flight (CyTOF) to monitor intracellular replication of *Salmonella enterica* serovar Typhimurium in a time-resolved animal model of infection. This technology was used to pinpoint specific host cell populations infected by minimal *Salmonella* variants expressing limited repertoires of Type 3 Secretion System (T3SS) effector genes. Mechanistically, these studies revealed a small subset of non-canonical monocytes which function as a primary cellular bottleneck to bacterial tissue transmission and the onset of Typhoid-like disease. We further determined how *Salmonella* overcomes this immunological bottleneck by expressing two effector gene networks, the θ effectors and the *spv* operon, that were acquired during distinct episodes of genomic expansion within the species. Importantly, the *spv* operon works in concert with the θ effector genes to overcome host cell-intrinsic immunity initiated by the Myd88 and IFN-γ signaling pathways. Our data further reveal SpvB as the critical member of the *spv* operon responsible for *S*.Tm colonization of lymphoid tissues. In this way, genome minimization has allowed us to define key evolutionary adaptations enabling food-borne systemic infection by *Salmonella* Typhimurium (Fig. 8b).

Several different infection models have been used to study the progression of *Salmonella* infection within the host organism. These models typically include either oral or intragastric (i.g.) administration of the pathogen to investigate intestinal mucosal infection, or intraperitoneal (i.p.) or intravenous (i.v.) administration to study mechanisms of systemic infection. Here, we chose to examine the tissue colonization dynamics of *S.*Tm at single cell resolution after i.g. delivery to track the small numbers of bacteria that bypass the intestinal mucosa and seed a secondary lymphoid organ (spleen) in a model that most closely mimics the natural route of infection ^52,53^. Importantly, our single cell analysis of *S.*Tm transmission within the spleen is consistent with many of the observations made using previous systemic infection models. For example, Geddes et al. reported *Salmonella* actively secreting SPI-2 effector proteins within monocytes, neutrophils, and B and T lymphocytes after i.p. injection of the pathogen^31^. Our mass cytometry analysis of LPS^pos^ signal largely recapitulated these observations. Surprisingly however, Geddes et al. did not identify infected mature macrophage (CD11b^pos^ F4/80^pos^), likely due to the very low numbers of detectable cells infected by *S.*Tm (Fig. 3d). Recently, Hoffman et al. reported that *S.*Tm can occupy two distinct macrophage populations identified as iNOS macrophage (CD11b^pos^ F4/80^pos^ Ly6C^pos^ CD69^pos^ MHCII^neg^) and CD9^pos^ macrophage (CD11b^pos^ F4/80^pos^ Ly6c^neg^ CD69^pos^ MHCII^neg^) in the spleen^54^. Our mass cytometry analysis of LPS^pos^ cells also revealed discrete populations of CD11b^pos^ F4/80^pos^ macrophage infected with *S.*Tm after i.g. inoculation, albeit at much lower levels than what was observed using the i.p. model of infection^54^.

While our mass cytometry data confirmed seminal observations in the field, it further extended our understanding of *S.*Tm infection dynamics by revealing a time resolved single cell atlas of cell-to-cell dissemination in a secondary lymphoid organ. Unexpectedly, we identified a CD62L^pos^ circulating monocyte population (sp22; CD11b^pos^ F4/80^neg^ CD62L^pos^) as an important cellular barrier to *S*.Tm infection of the spleen^23,28^. CD62L^pos^ monocytes were the earliest cell type infected with wildtype *S.*Tm and *S.*Tm θ *spv^+^* after i.g. inoculation. In addition, neither the *S.*Tm Efl strain that is unable to replicate intracellularly nor the *S*.Tm θ strain that is incapable of suppressing cell-intrinsic immune responses disseminated far beyond the CD62L^pos^ monocyte population. Interestingly, the progenitor CD62L^pos^ monocytes were no longer detected in middle or late-stage of either wild-type *S.*Tm or *S.*Tm θ *spv*^+^ infection. In these infections, loss of CD62L^pos^ monocytes correlated with the appearance of *S.*Tm infection of sp8 which shares markers consistent with a classically activated mature macrophage subpopulation (sp8 cells, CD11b^pos^ F4/80^pos^ MHCII^pos^). Notably, loss of the CD62L^pos^ monocyte population and dissemination of *S*.Tm into macrophage, neutrophils, B-cells, and T-cells only occurred in mice infected with S.Tm strains expressing the *spv* operon. It is currently unclear if this cell type undergoes cell death, resulting in release of *S.*Tm for uptake in new cells, or if these monocytes differentiate into activated macrophage. Previous studies have shown that immune cell death and efferocytosis is a key contributor to cell-to-cell transmission of *S.*Tm^21,55–57^. It is therefore attractive to speculate that *spv* operon genes promote cell death to aid in bacterial dissemination throughout the spleen. Interestingly, our analysis of the *spv* operon in tissue infection reveals SpvB as the critical effector responsible for *S*.Tm θ transmission. In addition, the SpvB effector protein has been implicated in macrophage cell death through ADP-ribosylation of G-actin^13,58^. Future studies will be needed to determine if and how the SpvB effector regulates CD62L^pos^ cells in the spleen and if *S*.Tm uses a common mode of cell-to-cell transmission in diverse lymphoid tissues.

It should be pointed out that the activated macrophage population (sp8) infected with *S*.Tm identified in our study do not appear to be the same cell type as the iNOS^pos^ or CD9^pos^ macrophage reported previously as early targets of the pathogen^54^. Some subsets of classically activated mature macrophage exhibit high levels of MHC class II surface expression whereas iNOS^pos^ and CD9^pos^ macrophage do not express this marker^54,59^. The discrepancies in these data likely result from different levels of infection using the two delivery methods. For example, *S.*Tm colonization of the sp8 macrophage population after i.g. delivery was observed at very low bacterial burdens (10^4^-10^5^) in our study. In contrast, infection of iNOS^pos^ and CD9^pos^ macrophage after i.v. inoculation of *S.*Tm were observed at two orders of magnitude larger spleen colonization (10^6^- 10^7^) ^54^. These data suggest that initial *S.*Tm seeding of the spleen after oral infection may be different from that observed after introduction of supraphysiological bacterial loads. However, our data does not conflict with the observation that iNOS^pos^ and CD9^pos^ are key cell types in *Salmonella* tissue colonization. As *S.*Tm replication reached the levels that are typically achieved by i.p. or i.v. infection, we observed *S.*Tm within F4/80^pos^ MHCII^neg^ macrophage (sp7) that may represent the iNOS^pos^ or CD9^pos^ macrophage previously reported^54^. Thus, late-stage progression of *S.*Tm colonization in the spleen after either oral or systemic infection likely occurs through a common mechanism.

The *Salmonella enterica* species is made up of over 2,500 serotypes, many of which have distinct host tropisms. Differences in the total number and composition of SPI-2 effector genes is a distinguishing feature of strains within the species. Remarkably, the θ effector genes are highly conserved across *Salmonella* serotypes, approximately 100 of which cause human disease^12^. It therefore appears that θ effector genes were acquired prior to the explosion of *Salmonella* taxonomic diversification. Furthermore, the θ genes have been retained in *Salmonella* species that have undergone episodes of genomic retraction, such as *Salmonella enterica* serovar Typhi. Thus, our studies indicate that θ effector genes provide the core molecular machinery for *Salmonella* spp. to replicate within a protective vacuolar environment of eukaryotic cells. We now propose that *Salmonella enterica* speciation has been driven in large part by coupling the core function of θ effector genes with novel evolutionary innovations that open new ecological niches to the pathogen. The studies presented here provide a glimpse into this process. Remarkably, whereas the *S.*Tm θ strain is unable to colonize lymphoid tissues, the host barriers to infection by this strain are broken in both *Myd88*^-/-^ and *Ifngr1*^-/-^ mice, or through co-expression of the *spv* operon effector SpvB. Thus, the acquisition of the *spv* effector genes was a key innovation in the evolution of serovar Typhimurium, allowing the pathogen to overcome host immunological barriers and to access new cellular habitats within deep tissues of susceptible mice.

A potentially controversial issue raised by our work is that many SPI-2 effector proteins have been implicated in the systemic colonization of *S.*Tm whereas our studies indicate that effector genes encoded by the minimal *S.*Tm θ *spv*^+^ strain are sufficient to drive the trajectory of cell-to-cell dissemination observed for wild type *S.*Tm. It should be pointed out however that our studies using the C57Bl/6 mouse model of infection are not exhaustive. We have not varied the initial inoculum nor did we examine the rates of tissue colonization at the earliest time points of infection. In addition, the *S.*Tm θ *spv*^+^ strain exhibits a very slight attenuation in virulence and the time to pathogenesis. Thus, additional effector proteins do contribute to the systemic disease model.

Finally, though our current findings are based on an oral route of *S*.Tm infection which more closely models the natural route of infection, these studies are restricted to the acute mouse model of infection. Further studies will be needed to corroborate these findings in bacterial persistence models. While the mass cytometry analysis was completed using a broad antibody panel, there is room for increasing the size and complexity of the antibody panel to further define the data set. This would allow detailed partitioning of the macrophage/monocyte subsets and better delineation of lymphocyte subtypes. Lastly, these studies were completed with *Salmonella enterica* in an SL1344 background. Though this is a canonical strain used in many studies, further experiments with other strains of *S.* Tm must be completed to confirm our findings in additional *Salmonella enterica* serovars.

## Extended Data

**Extended Data Fig. 1:**
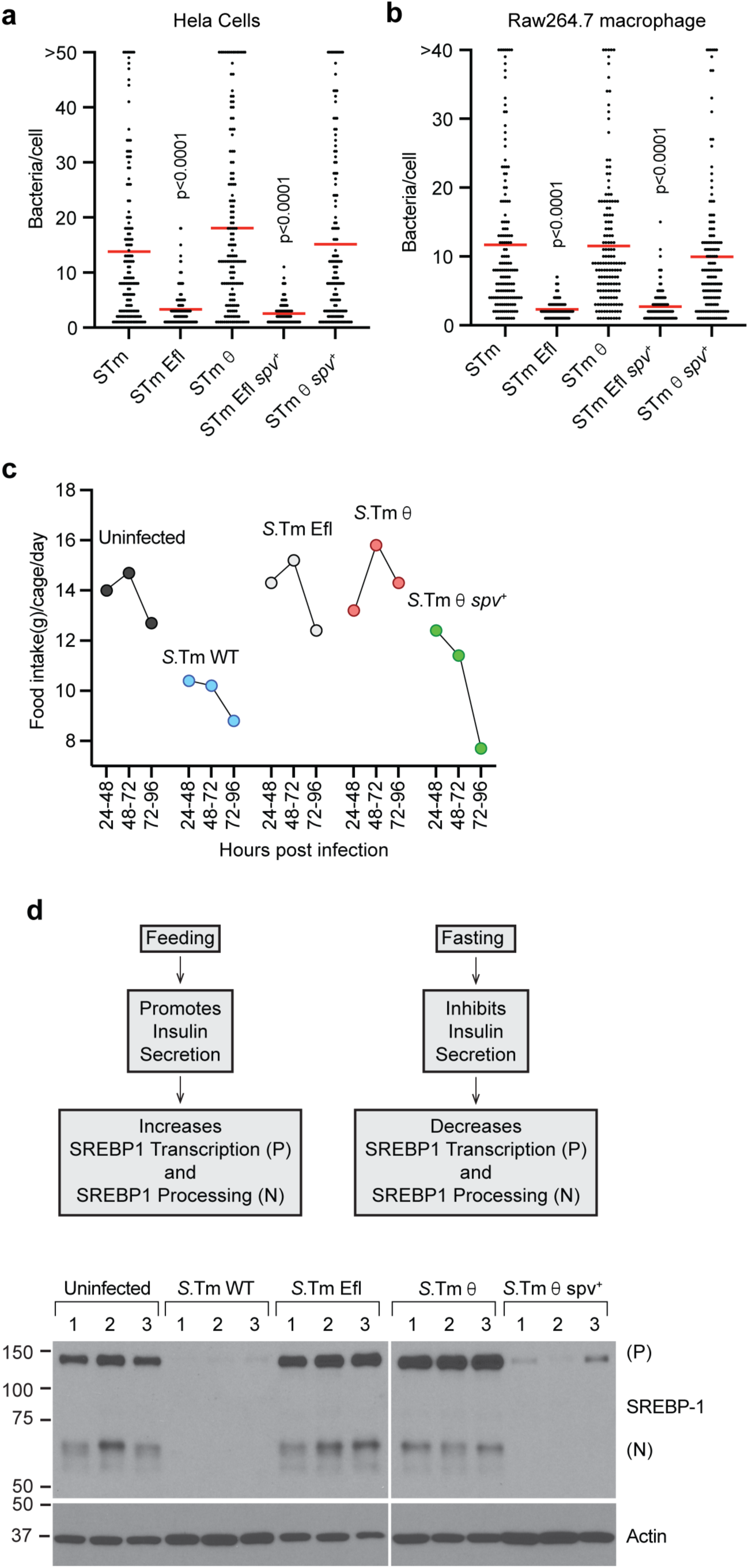
Characterization of *S*.Tm minimal effector strains. (a-b) Quantification of the number of individual bacteria per HeLa cell (a) or RAW 264.7 macrophage (b) infected with the indicated *S*.Tm strains. A total 150 cells from 3 biological replicates were counted. Infections were conducted on Hela (MOI = 50) and Raw 264.7 (MOI=1) and the number of bacteria per cell were quantified 18 hours post infection (hpi) and 24 hpi, respectively. (c) Food intake by C57BL/6J mice uninfected or infected intragastrically (i.g.) with 1×10^9^ *S.*Tm, *S.*Tm Efl, *S.*Tm θ, and *S.*Tm θ *spv*^+^ (n=5 mice/*S*.Tm strain). Food intake was measured at three different time points during a four day infection 24-48h, 48-72h, 72-96h. Food intake measurements for each cohort of mice included the entire cage (n=5 mice/cage). (d) Model of SREBP1 activation during feeding and fasting conditions. During feeding SREBP1 is processed and its N-terminal transcription factor domain translocates to the nucleus to initiate lipogenesis. During fasting SREBP1 levels decline. Western blots against SREBP1 showing full length unprocessed SREBP1 (P) or processed Nuclear (N) SREBP1 in livers from mice infected intragastrically (i.g.) with the *S.*Tm strains indicated. *S*.Tm strains used to infect mice (i.g.) with 1×10^9^ bacteria from the strains represented are shown at the top and each lane represents one mouse (n=3 mice for each *S*.Tm genotype).

**Extended Data Fig. 2:**
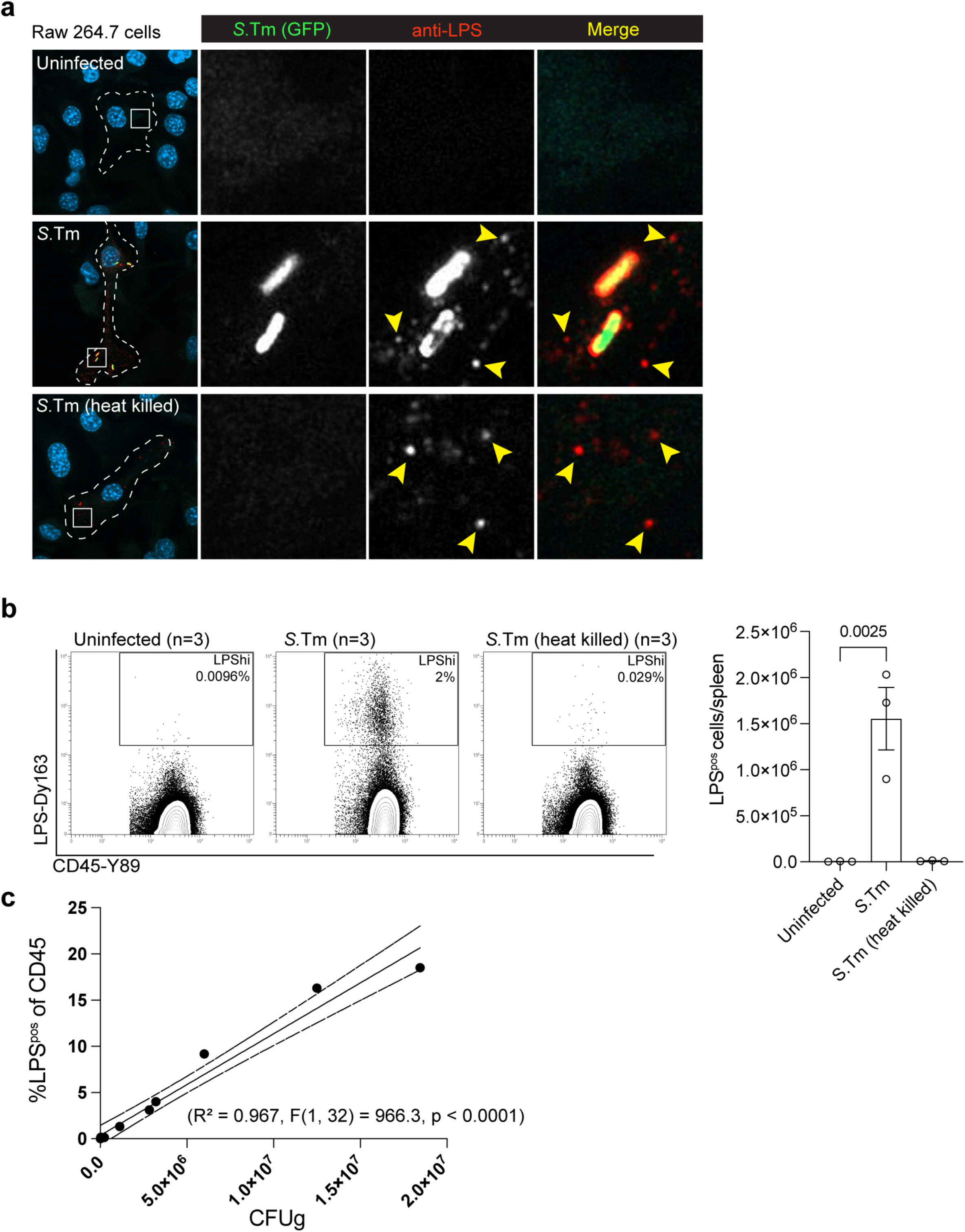
LPS staining with anti-LPS antibody *in vitro* and *in vivo*. (a) RAW 264.7 murine macrophages uninfected or infected with *S.*Tm-GFP or heat killed *S.*Tm-GFP (MOI = 100) and imaged 24 hpi and immunolabeled with anti-LPS antibody and stained with DAPI. Images are maximum intensity projections of confocal images spanning the entire z-dimension of the cell. Dotted lines show the cell border. Right panels are higher magnification images of the white boxed regions in left panels. (b) Flow Cytometry plots of spleens from C57BL/6 mice infected intraperitoneally (i.p.) with PBS (n=3), or 1×10^4^ *S*.Tm (n=3) or heat killed *S*.Tm (n=3) and stained with anti-*S*.Tm LPS-Dy^163^ and anti-CD45-Y^89^ antibodies (left). LPS^pos^ cells/spleen from the adjacent flow plots. (c) LPS^pos^ cells in mouse splenocytes from C56BL/6 mice (n=11) infected i.g. with *S*.Tm correlated with the CFU/g in spleen from the same mouse. Linear regression analysis performed on GraphPad Prism R^2^ = 0.967, F(1,32), p<0.0001.

**Extended Data Fig. 3:**
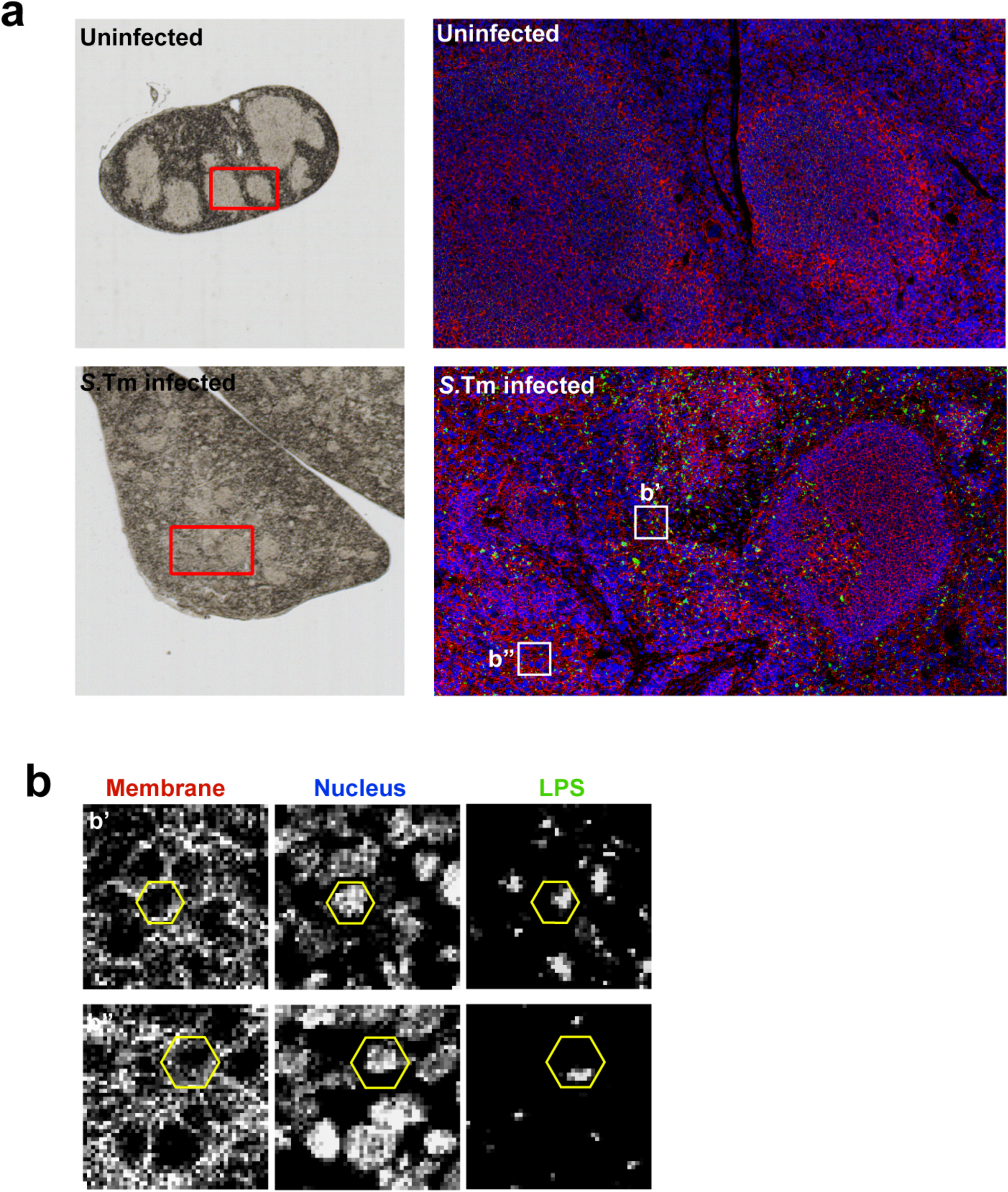
Imaging mass cytometry of *Salmonella* spleens. (a) Mouse spleen sections in bright field (left) and imaged with the Helios mass cytometer (right, higher magnification view of the red boxed region at (left) from uninfected (top) and i.g. infected (1×10^9^ *S.*Tm) (bottom). Sections labeled with DNA (blue), membrane marker (red), and *S.*Tm LPS (green). (b) (b’ – b’’) Insets from (a, right). Hexagons indicate the rough outline of individual cells identified by plasma membrane and nuclear labeling that are infected with *S*.Tm (identified by anti-LPS-Dy^163^).

**Extended Data Fig. 4:**
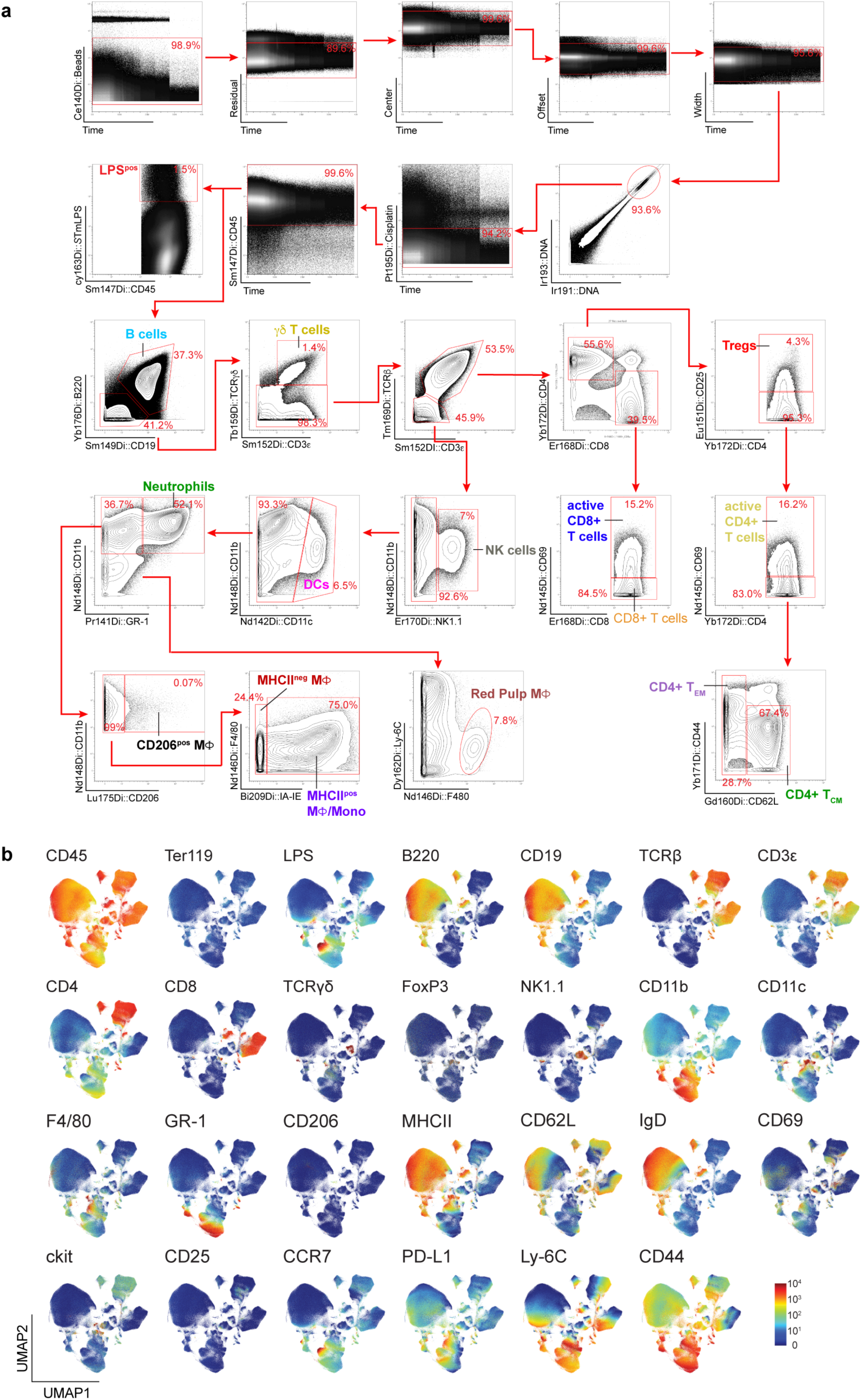
Gating strategy and UMAP signal distribution of immune markers for immune cell subsets. (a) Representative gating strategy for the 13 manually defined immune cell populations (see also Materials and Methods). Axes are labeled with the antibody epitope and the metal associated with the antibody. Red boxes are the gates used in the analysis pipeline to designate cells. Red arrows represent the next gate in the series. (b) UMAP Graphs showing heatmaps for the individual antibody markers in the immune panel. All graphs represent equal subsampling of each mouse (35,714 cells/mouse). Heatmaps include all 34 mouse spleens analyzed from all strains. The protein recognized by the antibody is labeled in the top left corner.

**Extended Data Table 1:** Data sets used in CyTOF analysis. Mouse samples used for CyTOF analysis including *S*.Tm strain used for infection, CFU/g, Log_10_CFU/g, days post infection the animal was sacrificed (dpi) and which figures include CyTOF data from this animal. All mice were infected intragastrically with 1×10^9^ of the indicated strains.

| Mouse number | Strain Genotype | Alto Lab Nomenclature | CFU/g | LogCFU/g | dpi | Figure 3 | Extended Data Figure 4 | Figure 4 | Figure 5 |
| --- | --- | --- | --- | --- | --- | --- | --- | --- | --- |
| 1 | uninfected | WB5-1 | 0 | 0.00 | 4 | a-g | a,b |  | a,b |
| 2 | uninfected | WB5-2 | 0 | 0.00 | 4 | a-g | a,b |  | a,b |
| 3 | uninfected | WB5-3 | 0 | 0.00 | 4 | a-g | a,b |  | a,b |
| 4 | S.Tm | WB1-9 | 0 | 0.00 | 4 | a-g | a,b | a |  |
| 5 | S.Tm | WB1-12 | 4,598 | 3.66 | 4 | a-g | a,b |  |  |
| 6 | S.Tm | WB1-3 | 10,390 | 4.02 | 4 | a-g | a,b |  |  |
| 7 | S.Tm | WB1-2 | 26,370 | 4.42 | 4 | a-g | a,b | a |  |
| 8 | S.Tm | WB1-8 | 231,707 | 5.36 | 5 | a-g | a,b | a |  |
| 9 | S.Tm | WB1-7 | 1,112,299 | 6.05 | 5 | a-g | a,b | a |  |
| 10 | S.Tm | WB1-10 | 2,826,087 | 6.45 | 4 | a-g | a,b | a |  |
| 11 | S.Tm | WB1-11 | 3,206,107 | 6.51 | 4 | a-g | a,b |  |  |
| 12 | S.Tm | WB1-4 | 5,992,509 | 6.78 | 7 | a-g | a,b |  | a,b |
| 13 | S.Tm | WB1-6 | 12,489,451 | 7.10 | 7 | a-g | a,b |  | a,b |
| 14 | S.Tm | WB1-5 | 18,449,612 | 7.27 | 7 | a-g | a,b |  | a,b |
| 15 | S.Tm Efl | WB4-1 | 0 | 0.00 | 4 | a-g | a,b |  |  |
| 16 | S.Tm Efl | WB4-2 | 0 | 0.00 | 4 | a-g | a,b |  |  |
| 17 | S.Tm Efl | WB4-3 | 0 | 0.00 | 4 | a-g | a,b |  |  |
| 18 | S.Tm Efl | WB4-4 | 0 | 0.00 | 7 | a-g | a,b |  | a,b |
| 19 | S.Tm Efl | WB4-5 | 0 | 0.00 | 7 | a-g | a,b |  | a,b |
| 20 | S.Tm Efl | WB4-6 | 0 | 0.00 | 7 | a-g | a,b |  | a,b |
| 21 | S.Tm $\theta$ | WB3-2 | 0 | 0.00 | 4 | a-g | a,b | b | |
| 22 | S.Tm $\theta$ | WB3-3 | 0 | 0.00 | 4 | a-g | a,b | | |
| 23 | S.Tm $\theta$ | WB3-4 | 0 | 0.00 | 7 | a-g | a,b | | a,b |
| 24 | S.Tm $\theta$ | WB3-6 | 0 | 0.00 | 7 | a-g | a,b | b | a,b |
| 25 | S.Tm $\theta$ | WB3-5 | 476 | 2.68 | 7 | a-g | a,b | | a,b |
| 26 | S.Tm $\theta$ | WB3-7 | 3,182 | 3.50 | 9 | a-g | a,b | b | |
| 27 | S.Tm $\theta$ | WB3-1 | 7,381 | 3.87 | 4 | a-g | a,b | b | |
| 28 | S.Tm $\theta$ | WB3-8 | 81,553 | 4.91 | 9 | a-g | a,b | b | |
| 29 | S.Tm $\theta$ spv+ | WB2-2 | 488 | 2.69 | 4 | a-g | a,b | c | |
| 30 | S.Tm $\theta$ spv+ | WB2-3 | 37,980 | 4.58 | 4 | a-g | a,b | c | |
| 31 | S.Tm $\theta$ spv+ | WB2-5 | 37,209 | 4.57 | 7 | a-g | a,b | c | a,b |
| 32 | S.Tm $\theta$ spv+ | WB2-1 | 315,232 | 5.50 | 4 | a-g | a,b | | |
| 33 | S.Tm $\theta$ spv+ | WB2-4 | 1,560,538 | 6.19 | 7 | a-g | a,b | c | a,b |
| 34 | S.Tm $\theta$ spv+ | WB2-6 | 2,061,433 | 6.31 | 7 | a-g | a,b | c | a,b |

## Materials and Methods

### Bacterial strains and growth conditions

This work was done using *S.*Tm SL1344 strain and its isogenic mutant derivatives. *Escherichia coli* DH5α was used for the construction and amplification of plasmids unless otherwise specified. Bacteria were routinely grown in Lennox L. Broth base (LB; Fisher), and, where appropriate, growth medium was supplemented with the following antibiotics at the indicated concentrations: ampicillin (100 μg/ml; Fisher), kanamycin (50 μg/ml; Sigma), streptomycin (100 μg /ml; Sigma), and chloramphenicol (15 μg/ml; Sigma). *Salmonella* strains expressing green fluorescent protein (GFP) were constructed by electroporation of the bacteria with the pBBR1MCS 6Y GFP plasmid^60^.

### Mammalian cell culture and *S*.Tm infections

Hela cells were grown in Dulbecco’s Modified Eagle Medium (DMEM) (Gibco, Thermo Fisher Scientific, supplemented with 10% Fetal Bovine Serum (FBS; bioWest) at 37°C in 5% CO_2_. RAW264.7 cells were grown in Roswel Park Memorial Institute (RPMI) Media (Gibco, Thermo Fisher Scientific, supplemented with 10% FBS (bioWest) at 37°C in 5% CO_2._ For microscopy, cells were seeded on coverslips in 6-well plates at a density of 7.5 x 10^4^ per well 24h before infection. Bacteria were grown in LB broth containing appropriate antibiotics for pBBRMCS selection (chloramphenicol, 8.5 µg/ml) overnight at 37°C. Bacterial cultures were diluted 1:33 in 10 ml of fresh medium and incubated at 37°C for 2.5h. Bacteria were diluted in cell culture medium and added to Hela cells at a multiplicity of infection (MOI) of 50 and RAW264.7 cells were infected at an MOI of 1. Cells were then centrifuged for 5 min at room temperature (800xg) to facilitate bacterial adherence. Invasion was allowed to proceed for 25 min at 37°C in 5% CO_2_, followed by incubation of the cells with 100 μg/ml gentamicin (Quality Biological) for 1h in the same conditions. The concentration of gentamicin was subsequently reduced to 20μg/ml gentamicin and cells were incubated for 18h for Hela cells and 24h for RAW264.7 cells (37°C, 5% CO_2_).

### Super resolution microscopy

Cells were plated on glass coverslips and infected the next day with GFP-labeled *S*.Tm (pBBR1MCS-GFP) at MOI 50 (Hela) or MOI 1 (RAW 264.7). At 18 h p.i. (Hela) or 24 h p.i. (RAW 264.7) cells were washed with PBS (Gibco), fixed with 3.5% PFA (Thermo) for 20 min, washed 3 times with PBS and then incubated with block/permeabilization buffer (0.1% saponin, 2% BSA in PBS) for 30 min. Cells were incubated with primary antibody (Rb anti-Lamp1, abcam ab24170, 1:500) for 1h, washed 3 times with PBS, then incubated with secondary antibody (goat anti-Rb Alexa 594, Invitrogen A11037, 1:200) for 1 h, followed by DAPI (1:20,000) for 20 min. Cells were then washed 3 times with PBS and coverslips mounted onto slides using VectaShield (Vector Laboratories H-1900). Z-stack images encompassing all bacteria in the cell were captured using a Zeiss LSM 980 laser scanning confocal microscope with Airyscan 2 at super resolution (x,y,z) and post-processed using Airyscan 3D SR (Zen 3.8) with autofilter and standard settings. Maximum Intensity Projections or a single z-plane image are shown, as indicated.

### Quantitative analysis of intracellular bacteria

Cells were plated on glass coverslips and infected the next day with GFP-labeled S.Tm (pBBRMCS-GFP) at MOI 50 (Hela) or MOI 1 (RAW 264.7). At 18 h p.i. (Hela) or 24 h.p.i. (RAW 264.7) cells were washed with PBS, fixed with 3.5% PFA for 20 min, washed 3 times with PBS and then with DAPI (1:20,000) for 20 min. Cells were then washed 3 times with PBS and coverslips mounted onto slides using VectaShield (Vector Laboratories H-1900). The cells were stained with DAPI. After fixation and mounting as above, cells were imaged and bacteria in individual cells were quantified using a fluorescent microscope (ZEISS Observer.Z1 microscope) with a 63x objective by starting at a fixed point on the slide and moving along the fields of view until bacteria within 50 cells were quantified. The observer was blinded to the experimental group during quantification. These experiments were repeated three times for a total of three infections (150 cells total) for each bacterial strain..

### Immunoblot for SREBP1

Harvested mouse liver tissue was snap-frozen in liquid nitrogen. 20mg frozen mouse liver was homogenized in tissue lysis buffer (1% SDS, 0.1% NP-40, 0.5mM EDTA, 1% TritonX-100, 5mM Tris-HCL and Complete Protease Inhibitor Cocktail Tablets [Roche]) in 2mL tough Microtubes with 2.8mm ceramic beads (Omni International), 4m/s 30s on 10s off for 3 rounds in a bead rotor (Omni 19010310). Samples were subject to centrifugation at (20,000xg, RT 10min) to remove debris. The protein concentration of the supernatant was determined by bicinchoninic acid assay kit (Thermo Scientific Pierce, 23227). 5x SDS sample buffer (BioRad) was added to samples then boiled at 55°C for 10 min and subjected to electrophoresis on 10% SDS-PAGE gels. The gel was transferred to a nitrocellulose membrane using Trans Blot Turbo (BioRad) and subjected to immunoblot staining for SREBP1 IgG-20B12 (5 μg/ml) and actin with anti-actin (1:1000; Cell Signaling-4970L). Bound antibodies were visualized using a 1:5000 dilution of goat anti-rabbit IgG (Jackson ImmunoResearch) conjugated to horseradish peroxidase and detection of the HRP signal using Supersignal West Pico PLUS Chemiluminescent substrate (Thermo Fisher). Membranes were exposed to Thermo Scientific CL-XPosure Film (Thermo Scientific) at room temperature for 10–120s.

### Staining samples for imaging mass cytometry

Serial paraffin sections were made for H&E histopathologic review and for Imaging Mass Cytometry. CyTOF slides were dewaxed and run to water before high intensity epitope retrieval in 10mM Tris/1mM EDTA pH 9 decloaking buffer held at near boiling temperature for 30 minutes via steam. Slides were cooled to room temperature, blocked with 3% bovine serum albumin, and incubated with the panel of metal conjugated primary antibodies listed in table below. Sections were stained with iridium DNA intercalator, washed, air dried and imaged for ROI mapping against their matching H&E slides. Dilutions given for antibodies from manufacturer (StandardBiotools) and optimal concentrations given for antibodies conjugated at UT Southwestern in table below. Data was acquired on the Hyperion Imaging System (Standard Biotools) and data processed with MCDViewer (Standard Biotools) and imported into Adobe Illustrator for downstream processing.

| Antibody Target | Clone | Metal | Species | Dilution or [Optimal] (µg/ml) | Product Number | Source |
| --- | --- | --- | --- | --- | --- | --- |
| SMA | 1A4 | 141Pr | Mouse | 1:100 | 2491903 | StandardBiotools |
| CD44 | IM7 | 144Nd | Mouse | 5µg/ml |  | UTSWClone |
| PanCytokeratin | AE1/AE3 | 148Nd | Mouse | 1:300 | 1361904 | StandardBiotools |
| CD11b | EPR1344 | 149Sm | Mouse | 1:300 | 2108703-11 | StandardBiotools |
| Fibronectin | EPR19241-46 | 152Sm | Mouse | 1:150 | 23115388-16 | StandardBiotools |
| FoxP3 | FJK-16S | 155Sm | Mouse | 1:50 | 2111860-12 | StandardBiotools |
| CD4 | BLR167J | 159Tb | Mouse | 1:100 | 2111865-12 | StandardBiotools |
| iNOS | SP126 | 160Gd | Mouse | 1:150 | 2212661-21 | StandardBiotools |
| B220 | RA3-6B2 | 162Dy | Mouse | .5µg/ml |  | UTSW Clone |
| S.Tm LPS | IE6 | 163Dy | Mouse | 1µg/ml | AB8272 | AbCam (This study) |
| F4/80 | D2S9R | 166Er | Mouse | 1:100 | 2111857-12 | StandardBiotools |
| Collagen1 | Polyclonal | 169Tm | Mouse | 1:300 | 281902 | StandardBiotools |
| GranzymeB | EPR22645-206 | 171Yb | Mouse | 1:250 | 2111861-12 | StandardBiotools |
| CD31 | EPR17259 | 172Yb | Mouse | 1:2000 | 2111854-12 | StandardBiotools |
| CD45 | D3F8Q | 175Lu | Mouse | 1:100 | 2111855-12 | StandardBiotools |
| CD8 | EPR21769 | 176Yb | Mouse | 1:100 | 2212407-05 | StandardBiotools |
| ICSK1 | Proprietary | 195Pt | Mouse | 1:100 | 201500 | StandardBiotools |
| ICSK2 | Proprietary | 196Pt | Mouse | 1:100 | 201500 | StandardBiotools |
| ICSK3 | Proprietary | 198Pt | Mouse | 1:100 | 201500 | StandardBiotools |

### *S.* Typhimurium infection studies in mice

All experiments involving mice were approved by the Institutional Animal Care and Use Committee at UT Southwestern Medical Center (APN# 2018-102566). 8-10 weeks old C57BL/6J (Jackson Laboratories 000664), *Ifngr1^-/-^* (Jackson Laboratories 003288), *Myd88^-/-^* (Jackson Laboratories 009088), and *Ifnar1^-/-^*(Jackson Laboratories 028288) bred and maintained under specific pathogen-free conditions in the animal care facility at UT Southwestern Medical Center were used for all experiments. Mice were infected as described below. Bacterial loads in the tissues of infected mice were evaluated at time points between 4 days post infection and 15 day survival. For *S.*Tm intraperitoneal (i.p.) infections, *S.*Tm strains were cultured to log-phase growth in LB broth in a shaker at 37°C, washed and diluted with sterile PBS, and injected intraperitoneally with a dose of 1×10^4^ CFUs per mouse. For *S.*Tm intragastric (i.g.) infection, *S.*Tm strains were grown overnight in LB broth supplemented with 100 μg/ml streptomycin (Sigma) in a shaker at 37°C and then diluted to 10^9^ CFUs in 0.1 ml LB. *S.*Tm strains were loaded in a feeding needle and delivered to mice by gavage. Bacterial loads in different tissues were determined by homogenizing organs with a Tissue homogenizer (OMNI) in 2 ml PBS and by plating serial dilutions on LB plates containing streptomycin. For survival experiments, mice were monitored daily for 15 days. Mice that exhibited a weight loss of greater than 20% of their starting weight or presenting with a cumulative clinical score of 4.5 according to UT Southwestern Animal Care Facility protocols were euthanized via carbon dioxide followed by cervical dislocation.

### Conjugation of LPS antibody for mass cytometry

Anti-Salmonella LPS antibody (abcam ab8274) was conjugated to 163Dy Lanthanide metal with the MaxparX8 metal conjugation kid (StandardBiotools Cat# 201163A) according to manufacturer’s instructions. In brief: Add 95μl L-buffer to X8 Polymer with 50mM 163Dy chloride solution. Add 5μl 50mM metal solution to X8 polymer tube. Incubate 37°C 40min in water. In a separate tube add 100μg stock antibody to 50kDa filter and adjust volume to 400μl with R-buffer. Centrifuge (12,000xg, RT, 10min). After polymer incubation add 200μl L-buffer to 3kDa filter and centrifuge at (12,000xg, RT, 25min). Add 100μl TCEP solution to 50kDa antibody filter and immediately incubate at 37°C for 30min. Wash x8 polymer in 3kDa filter in 400μl C-buffer and centrifuge at 12,000xg 30min. Wash 50 kDa filter unit with antibody twice 400μl C-buffer and spin (12,000xg, RT, 10min). Resuspend metal loaded polymer in 60μl C-buffer and transfer contents to 50 kDa filter with antibody and mix gently. Incubate at 37°C for 90min in a water bath. After conjugation is complete add 200μl W-buffer to the 50kDa filter with the 100μl total antibody conjugation mixture and centrifuge at (12,000xg, RT, 10min). Repeat wash steps with W-buffer three times. Resuspend in 80μl W-buffer and measure protein concentration using a NanoDrop. Add the necessary volume of Antibody Stabilizer PBS (StandardBiotools) to have a final concentration of 0.5mg/ml antibody. Spin 50kDa column upside down into collection tube at (1,000xg, RT, 2min). Store antibody at 4°C for future use.

### Staining samples for suspension mass cytometry

8-10 weeks old C57BL/6 mice were orally infected with 1×10^9^ CFUs of *S.*Tm strains as described above. To determine immune cell counts in the spleen, mice were sacrificed, and spleens were mechanically homogenized though a 70μM cell strainer (Falcon) with a 1ml Plastic syringe (BD Biosciences) into 2ml RPMI (Gibco) in a 50ml conical (basix). The homogenized filter was washed with 10mL RPMI buffer and passed through the filter two more times. Cells were pelleted (300xg, 4°C, 4min). Red blood cells were lysed with 5ml 1xRBC lysis buffer (Tonbo) and incubated at room temperature for 5min. Phosphate Buffer Saline (PBS)(Sigma) was added for a final volume of 20ml. Cells were pelleted and resuspended in 4ml PBS and transferred to a 15 ml conical (Falcon). 100ul of the cell suspension was used for plating bacterial CFU for the sample. Whole spleens were incubated in 1ml RPMI 5 µM CellID-Cisplatin (Standard Biotools) for 20min. After incubation add 5ml of Cell Staining Buffer (Standard Biotools). Cells were pelleted (300xg, 4°C, 4min) and resuspended in 4ml Cell staining buffer. Cells were pelleted (300xg, 4°C, 4min) and resuspended in 50ul 1:100 FcBlock (anti-CD16/CD32 (Tonbo)) 4°C, 10min. 50ul Cell Staining Buffer with antibodies for surface antigens (See table below) were added for 30 minutes and vortexed every 15min. Dilutions given for antibodies from manufacturer (StandardBiotools) and optimal concentrations given for antibodies conjugated at UT Southwestern in table below. Cells were washed and pelleted twice with 2ml Cell staining buffer. Pelleted cells were incubated in Fix I Buffer (Standard Biotools) for 15min at room temperature. Cells were washed in 2ml Maxpar Perm-S Buffer and pelleted (800xg,4°C, 4min). Cells were incubated in 2ml Maxpar Perm-S buffer for 20min then pelleted (800xg,4°C, 4min). All but 50ul of the supernatant was removed. 50ul *S*.Tm LPS-Dy^163^ was added and incubated for 30min. Cells were washed and pelleted twice (800xg,4°C, 4min) with 2ml Maxpar cell staining buffer. The pellet was then fixed with a 1.6%PFA solution (diluted in PBS form 16% PFA (Thermo) for 10min at room temperature. Cells were pelleted (800xg,4°C, 4min) and resuspended in 1ml MaxParFix and Perm Buffer with 0.125 µM Intercalator (Standard Biotools) overnight at 4°C. Cells were then washed and pelleted (800xg,4°C, 4min) in 2ml Cell staining buffer and then resuspended in 1mlCell Acquisition Solution (Standard Biotools) counted and run on the Helios Mass Cytometer System (Standard Biotools). Data was analyzed on the OMIQ software (Dotmatics).

| **Antibody Target** | **Clone** | **Metal** | **Dilution or [Optimal] (µg/ml)** | **Product Number** | **Source** |
| --- | --- | --- | --- | --- | --- |
| Ly6G/C/GR-1 | RB6-8C5 | 141Pr | 1:100 | 3141005C | StandardBiotools |
| Cd11c | N418 | 142Nd | 1:100 | S3142003C | StandardBiotools |
| CD69 | H1.2F3 | 145Nd | 1:100 | 3145005C | StandardBiotools |
| CD45 | 30-F11 | 147Sm | 1:100 | 3147003C | StandardBiotools |
| CD11b/Mac-1 | M1/70 | 148Nd | 1:100 | 3148003C | StandardBiotools |
| CD19 | 6D5 | 149Sm | 1:100 | 3149002C | StandardBiotools |
| CD25 | 3C7 | 151Eu | 1:100 | 3151007C | StandardBiotools |
| CD3e | 145-2C11 | 152Sm | 1:100 | 3152004C | StandardBiotools |
| TER-119 | TER-119 | 154Sm | 1:100 | 3154005C | StandardBiotools |
| CD62L | MEL-14 | 160Gd | 1:100 | 316008C | StandardBiotools |
| CD8a | 53-6.7 | 168Er | 1:100 | 3168003C | StandardBiotools |
| TCRBeta | H57-597 | 169Tm | 1:100 | 3169002C | StandardBiotools |
| NK1.1 | PK136 | 170Er | 1:100 | 3170002C | StandardBiotools |
| CD44 | IM7 | 171Yb | 1:100 | 2009641-14 | StandardBiotools |
| CD4 | RM4-5 | 172Yb | 1:100 | 3172003C | StandardBiotools |
| B220 | RA3-6B2 | 176Yb | 1:100 | 3176002C | StandardBiotools |
| IA/IE | 114.15.2 | 209Bi | 5µg/ml | 3209006B | UT Southwestern |
| Ly6C | HK1.4 | 162Dy | 5µg/ml | 3162014B | UT Southwestern |
| CCR7 | 4B12 | 164Dy | 5µg/ml | 3164013A | UT Southwestern |
| PDL-1 | 10F.9G2 | 153Eu | 5µg/ml | 3153016B | UT Southwestern |
| FoxP3 | FJK-16S | 158Gd | 5µg/ml | 3158003A | UT Southwestern |
| CD206 | C068C2 | 175Lu | 5µg/ml | 141702 | UT Southwestern |
| F480 | BM8 | 146Nd | 5µg/ml | 3146008B | UT Southwestern |
| IgD | 11-26c.2a | 150Nd | 5µg/ml | 3150011B | UT Southwestern |
| TCRgd | GL3 | 159Tb | 5µg/ml | 3159012B | UT Southwestern |
| CD117 | 2B8 | 173Yb | 5µg/ml | 3173004B | UT Southwestern |
| S.Tm LPS | 1E6 | 163Dy | 5µg/ml | AB8274 | AbCam(This study) |
| Fc Block |  |  | 5µg/ml | 70-0161-U500 | Tonbo |

### Mass Cytometry Data Analysis

All Mass Cytometry analysis was conducted using the OMIQ platform (Dotmatics). The analysis parameters are outlined and all data is accessible (see Data Availability Statement). The stained spleen samples (totaling 34 spleens) were gated according to the Gaussian parameters illustrated in Extended Data Fig 4a, as well as the bivariate plots used to distinguish canonical immune cells and their requisite immune cell markers in Extended Data Fig. 4a. Following this, the samples were evenly subsampled for analysis to 35,714 cells per sample from the CD45^pos^ gate, which had been filtered through the following gates: Beads, Residual, Center, Offset, Width, DNA (cells) Cisplatin (live/dead) and CD45^pos^. After that the cells were designated with canonical immune cell classifications represented in detail (Extended Data Fig. 4) Briefly the cell types were designated: B-cells (CD45^pos^ CD19^pos^ B220^pos^), γδT-cells (CD45^pos^ CD3ε^pos^ TCRγδ^pos^), Naïve CD4 T-cells(CD45^pos^ TCRβ^pos^ CD3ε^pos^ CD62L^pos^),T-regs(CD45^pos^ TCRβ^pos^ CD3ε^pos^ CD4^pos^ CD25^pos^),activeCD4T-cells (CD45^pos^ TCRβ^pos^ CD3ε^pos^ CD4^pos^ CD69^pos^), Eff/memCD4T-cells (CD45^pos^ TCRβ^pos^ CD3ε^pos^ CD4^pos^ CD62L^lo^) Central memory CD4T-cells (CD45^pos^ TCRβ^pos^ CD3ε^pos^ CD4^pos^ CD62L^high^), CD8 T-cells (CD45^pos^ TCRβ^pos^ CD3ε^pos^ CD8^pos^), active CD8 T-cells (CD45^pos^ TCRβ^pos^ CD3ε^pos^ CD8^pos^ CD69^pos^), NK Cells (CD45 ^pos^ NK1.1 ^pos^), Dendritic Cells (DC’s) (CD45^pos^ CD11c^pos^), Neutrophils (CD45^pos^ CD11b^pos^ GR-1^pos^), CD206^pos^ Macrophage (Mϕ) (CD45^pos^ CD11b^pos^ CD206^pos^) MHCII^neg^ Mϕ (CD45^pos^ CD11b^pos^), MHCII^pos^ Mϕ (CD45^pos^ CD11b^pos^ F4/80^pos^ IA/IE^pos^) Red Pulp Macrophage (CD45^pos^ CD11b^pos^ Ly6C^pos^ F4/80^pos^).

Dimensionality reduction was performed using UMAP with the following settings: Neighbors = 15; Minimum Distance = 0.4; Metric: Euclidian; Learning Rate: 1; Epochs = 200; Random Seed = 3641; Embedding Initialization = Spectral. Collected LPS^pos^ cells were subjected to UMAP (Neighbors = 15; Minimum Distance = 0.4; Metric: Euclidian; Learning Rate: 1; Epochs = 200; Random Seed = 3641; Embedding Initialization = Spectral) and PhenoGraph analysis using K-value = 20; Distance Metric: Euclidian; Louvain Seed: 2126. These parameters were used to generate heatmaps, cell counts, and values for downstream cellular data analysis.

To determine the number of cells in each population, the total number of immune cells per sample were determined by back-calculating the percentage of cells of interest as a proportion of the CD45^pos^ gate. This percentage was then multiplied by the total number of live cells in the full spleen samples after red blood cell lysis (as described above). A full table of data used for these calculations is available (see our Data Availability Statement).

### Statistical analysis

Unless otherwise indicated all statistical analysis was completed using GraphPad9 (Mathworks). Mass Cytometry experimental data processing and analysis for: UMAP, PhenoGraph, and FlowSom analysis were all completed in the OMIQ software (Dotmatics). All experiments where applicable were performed at least three times independently. A p-value of less than 0.05 was considered significant.

## Acknowledgements

We would like to give special thanks to Sebastian Winter and the Winter lab (UC Davis) for their assistance in developing our bacterial strains and mouse experiments used in this experiment. We would further like to acknowledge the UT Southwestern Microbiology Department Live Cell Imaging Facility (NIH: 1S10OD034383) for the use of their microscope and technical expertise, We would also like to acknowledge Angela Mobley and the other members of the UT Southwestern Flow Cytometry Core Facility for their assistance with mass-cytometry. Furthermore, we would like to mention the UT Southwestern Histopathology Core including John Shelton and Alejandro Daniel for their histopathology and imaging mass-cytometry staining. This work was funded by the National Institute of Health (AI083359 to NMA; AI158357 to AR and NMA), The Welch Foundation (I-1704 to NMA and I-1793 to AR), and The Burroughs Welcome Fund (1011019 to NMA).

## Data availability statement

The data generated in this study has been deposited in the Open Science Forum database (https://osf.io) in accordance with the requirements of Nature Microbiology. All FCS files, along with detailed sample annotations and panel information (including metal tag), are available within the repository.

Preprocessing and gating strategies were performed using OMIQ data platform (https://www.omiq.ai) and are described in Supplementary Methods. The high-dimensional clustering and downstream analyses (e.g., UMAP, PhenoGraph) were performed using the OMIQ platform and the data parameters available in the Materials and methods.

Further information and requests for raw data or materials should be directed to the corresponding author.

## Inclusion and Ethics Statement

This study was conducted in accordance with the ethical standards of the University of Texas Southwestern Medical Center and the relevant national and international guidelines. All necessary ethical approvals were obtained prior to data collection.

## Author Contributions

N.M.A. and W.B.B. designed the study and wrote the manuscript. W.B.B., L.T.A., H.D., D.M., and A.B.M. performed the experiments and participated in the data analysis. J.D.F. assisted in the mass-cytometry experiment design and data analysis. A.R. assisted in the metabolic activation experiment design and data analysis.

